# Scale dependent temporal and spatial patterns of population change in anadromous sea trout (*Salmo trutta*) in Scotland over seven decades

**DOI:** 10.1101/2025.11.07.687197

**Authors:** J.A. Dodd, I.E. Moore, Adrian W. Bowman, C.W. Bean, J.R. Rodger, C.E. Adams

**Author notes:** Corresponding authors: Jennifer A. Dodd, Centre for Conservation and Restoration Science, School of Applied Sciences, Edinburgh Napier University, Sighthill Campus, Edinburgh, EH11 4BN, Scotland. Funder information- Funding for this research was provided by Grieg Seafood Ltd and the Skye and Lochalsh Rivers Trust (formerly part of the Skye and Wester Ross Fisheries Trust).

## Abstract

The spatial scale at which temporal change in a population is measured has the potential to skew the inferences drawn on patterns of change from monitoring data. This study analysed data from rod catches of anadromous sea trout (*Salmo trutta*) in Scotland over a period of 72 years and over three spatial scales (National, Regional and District). The pattern of temporal change evident at the national scale mostly did not clearly manifest at Regional or at District spatial scales; which were also generally decoupled (67% of Regions and 80% of Districts exhibited a different pattern from the National pattern). At the National level, very clear and substantive declines over the study period were evident. At Regional levels (N=9) change in rod catches was more nuanced with some Regions showing no change, one Region showing an increase in sea trout caught and 6 (67%) exhibited a different pattern from the National one. At the District levels (N=64) opposing patterns of change were observed over the study period, even in adjacent Districts with 80% of Districts exhibiting a different pattern from the National one. A conclusion of this study is that, as a national resource, sea trout numbers have declined substantively over the last seven decades (by around 50%). However, this general picture is not applicable across all geographic Regions nor all Districts. Two logical inferences derive from these results. Firstly, for best effect, the application of limited resources for management should be focused at smaller scales than at countrywide. Secondly, that monitoring population change at small spatial scales, for example in several index rivers, is unlikely to capture the highly nuanced patterns of change in species with high levels of intra-specific structuring as is commonly exhibited by the Salmonidae.

## INTRODUCTION

The spatial and temporal scales over which ecological processes are investigated can strongly influence observed patterns in the resultant data and thus their subsequent interpretation (Levin, 1992; Wiens, 1989). For example, Johnson and colleagues (Johnson *et al*., 2004) showed marked differences in the spatial scale at which three Arctic mammal species respond to the available vegetation resources. Similarly, Sherry and Holmes (1988) show that at broad spatial scales, there is a strong positive correlation between the presence of two forest birds, the Least Flycatcher, *Empidonax minimus* and the American redstart, *Setophaga ruticilla*, suggesting ecological co-existence. At smaller spatial scales however, the former species had a marked negative effect on the latter which was manifest though strong competition for breeding sites (Sherry & Holmes, 1988). For both these examples, and many others in an expanding literature on this topic, studies on the same systems but at different spatial scales could lead to manifestly different inferences about the ecological processes under scrutiny (Wiens 1989).

One applied use of ecological data through which scaling might lead to inappropriate conclusions, is in the assessment of the status of a wildlife feature of interest. For example Murdoch & Aronson, (1999) showed that the spatial scale over which data on coral reef cover was collected had the potential to significantly impact the assessment of their conservation status. For most species, an assessment of their conservation value is made at relatively broad spatial scales. Most frequently the scale of assessment is defined by political and jurisdictional boundaries; frequently country but occasionally state or province. Many nations use legislative tools to identify important species for conservation. For example, in the United States, this is done through the Endangered Species Act (1973); in South Africa, through the National Environmental Management: Biodiversity Act (2004); and in Australia, through the Environmental Protection and Biodiversity Conservation Act (1999). Although it is implicit in this process that, having identified priority species, resources will be allocated towards delivery, in reality, it is unlikely that all priority species will also be prioritised for funding.

One of the key concerns about priority species lists is that assessment of the status of a species at the level of country may not be the most appropriate geographic scale to provide meaningful insights. There is good reason to argue that this might be particularly the case for species that exhibit some form of intraspecific structuring (Koene *et al*., 2024). This occurs in species that occupy habitats that are spatially disjunct, where the species has poor dispersal ability and/or where gene flow across the species is constrained either by geography or by the behaviour of the species (Bush & Adams, 2007). This is relatively common in species that inhabit freshwater ecosystems (Koene *et al*., 2024) where dispersal between discrete populations can be severely restricted (Maitland & Adams, 2001) and amongst migratory species which exhibit strong natal homing leading to distinct populations (Adams *et al*., 2006; Gilbey *et al*., 2017; Rodger *et al*., 2024). One migratory fish species that exhibits strong intraspecific structuring is the brown trout, *Salmo trutta.* The species is widely distributed across the northern hemisphere and has been introduced to many parts of the southern hemisphere (Ferguson *et al*., 2019a). Many populations are partial migrants with a component of the population remaining in freshwater throughout their life and another component migrating to sea; this anadromous component is hereafter referred to by the common name for this life history strategy of this species, ‘sea trout’, (Ferguson, 2006).

Despite that across much of its range, sea trout support an important recreational and, in places, commercial fishery (Lobón-Cerviá, 2009; ICES, 2016; King *et al*., 2016; 2024) reliable assessments of population status, long-term change in populations and inter-annual population variation are rare (ICES, 2013; Höjesjö *et al*., 2017; Wilson & Veneranta, 2019). The few existing assessments indicate that the size of European sea trout populations have changed over time. In the Baltic Sea, England and the Netherlands, some sea trout populations appear to have declined since the 1990’s (ICES 2013; Harris & Evans, 2017). In contrast, other countries have seen an increase in numbers. Denmark and Sweden, for example, reported a threefold increase in their sea trout populations between the 1980’s and 2010’s (ICES, 2013; Höjesjö *et al*., 2017). A national assessment of the status of sea trout in Norway showed that in only 25% of rivers were sea trout assessed as being of good or very good status (Fiske *et al*., 2024). This suggests that changes in population abundance are not uniform across the species’ range and that both the drivers of change and their relative influence likely vary across spatial scales.

The exact number of sea trout populations in Scotland is unknown, this is in part because of their ability to establish in small, short coastal river systems and thus may remain unrecorded. Defining the boundaries of a sea trout population is also problematic. The anadromous (sea trout) component of a population of *Salmo trutta* most frequently comprises a sub-set of individuals drawn from a wider gene pool that also comprises individuals that reach maturity without migrating to sea (frequently referred to as resident brown trout) (Ferguson et al. 2019b; Rodger et al. 2021). Both anadromous and non-anadromous components can exhibit high levels of genetic structuring. Thus, fish from different streams in the same river catchment may have highly restricted gene flow and thus may reasonably be regarded as discrete populations (Ferguson, 1989; Rodger et al. 2024). Because of the difficulty of defining population boundaries, here we use the term population in the looser sense of: individuals with the potential to interbreed living in the same area (see Berryman, 2002). Despite this, it is clear that sea trout are widespread across Scotland and are an important national fishery resource (Lamond, 1916; Menzies, 1936; Milner *et al*., 2006; Nall, 1930). Published studies examining temporal patterns of change in sea trout populations in Scotland, are limited to short time-series and geographically constrained areas, such as specific catchments or Regions (e.g. Pratten & Shearer, 1985; Shelton, 1993; Butler & Walker, 2006; Davidson *et al*., 2017; Adams *et al*., 2022). For example, Pratten & Shearer (1985) reported that sea trout captured in commercial netting stations in the North Esk, a large east coast river catchment in Scotland which drains into the North Sea, exhibited a 10–15 year cyclical pattern in abundance. As with the broader geographic pattern, the limited studies of temporal trends in sea trout catches hint at geographically contrasting patterns of change. Shelton (1993) and Butler & Walker (2006) reported long term declines in sea trout populations from the north west of Scotland, whereas recent increases in population size are indicated by catches in the north east (Davidson *et al*., 2017).

To-date there has been no systematic national analysis of the change in sea trout populations across Scotland. Since 1952, both commercial and recreational fisheries in Scotland have been required by law to report annual catches of Atlantic salmon and sea trout to the Scottish Government (Scottish Government 2024a).

Here we use this dataset to examine both the spatial and temporal variation in sea trout catches in Scotland. Specifically, we:

1. describe and contrast the change in sea trout catch size in Scotland between 1952 and 2023 at three spatial scales: National, Regional and District;
2. identify periods of statistically significant change in catch size at the different spatial scales.

## MATERIALS AND METHODS

### Data sources

The “Scottish Salmon and Sea Trout Fishery Statistics” (hereafter the SSSTF dataset), is a large, publicly available dataset (Scottish Government, Marine Directorate, 2024a), comprising a compilation of the reported numbers of Atlantic salmon (*Salmo salar*) and sea trout captured by three different methods: two primarily commercial: coastal and estuarine methods (“fixed engine” and “net and coble” fishing) and one recreational (“rod”). The data are collected as catch returns from fishery managers and owners and compiled by “District” which is a spatial scale that comprises either a single river catchment or a number of adjacent river catchments (Table S1). In 1994, a modification to the method of capture information resulted in rod caught fish being recorded as either “Rod: Retained” (where fish were removed from the watercourse) and “Rod: Released” (where fish were returned to the watercourse after capture). In 2004, an additional category was added for captured “finnock” (sea trout weighing less than 0.5kg; Marine Scotland, 2015).

### Data quality control and improvement

Due to declining fishing activity and spatial inconsistencies in commercial fishing for sea trout, these data were not included in the analyses presented here (Youngson *et al*., 2002). Thus, only recreational angling rod catches from between 1952 and 2023 inclusive were analysed in this study.

The SSSTF dataset does not provide any measure of fishing effort for the rod fishery (Marine Scotland, 2015). However, rod catch data, uncorrected for effort, has been shown to be a good proxy for population size in a number of studies. With several studies showing strong relationships between rod catch data and the number of fish detected by fish counters, as well as similar trends between raw catch data and catch data that has been corrected for fishing effort (see Youngson *et al*. 2002; Beaumont *et al*. 1991; Crozier & Kennedy 2001; Adams *et al.,* 2022; Thorley *et al*. 2005 for examples from Atlantic salmon and Smith *et al*. 2000 for examples from steelhead trout (*Oncorhynchus mykiss*)). An underlying premise of the study presented here is that trends in rod catch data for sea trout derived from the SSSTF database, uncorrected for fishing effort can provide insights into spatial and temporal abundance trends for this species in Scotland.

For the purposes of this study, annual reported catches of sea trout are defined as, sea trout and finnock captured by rod fishing (including both retained and released) totalled for each year separately. The SSSFT database reports catches from 109 Districts, from which 106 were identified that reported rod catch data for sea trout. To improve data quality and reduce false-zero data (Grueber *et al*., 2011), Districts which only provided incomplete catch records were excluded from further analysis; 30 Districts were thus removed from further analysis. Zero entries in the SSSTF dataset are generated from two different processes: a catch return where no sea trout were captured in that year (a true zero) and where there was no catch return made for that year (which could be either a true zero or a false zero (no data)). Unfortunately, it is not possible to distinguish between these two zero data types (Marine Scotland, 2015). For this reason, Districts with records comprising 25% or more zeros across all years were removed from further analysis. A further twelve catchments were removed on this basis. Thus, the resulting dataset used for further analysis comprised data from 64 Districts (Table S1).

### Spatial Scale

To test for the effect of the spatial scale on the trends in sea trout catches, models describing temporal change were constructed at three spatial scales. These were at a National (whole of Scotland, excluding the Orkney and Shetland Island groups), Regional (n=9; Figure 1a) and District scales (n=64; Figure 1b; Table S1).

**Figure 1.**
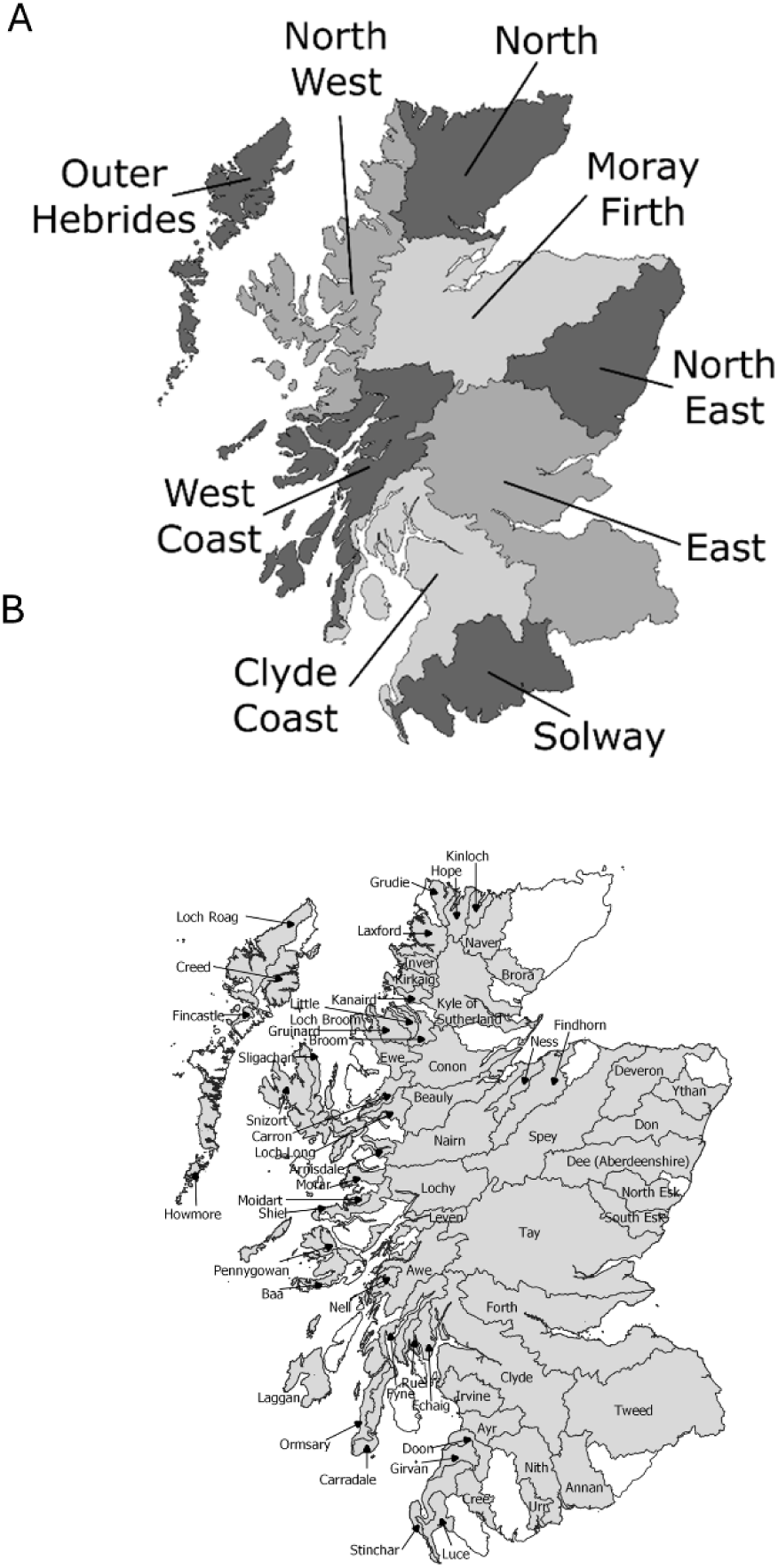
a) The nine Regions comprising b) the 64 Districts (see Table S1) across Scotland which were included in this study.

### Ethical Statement

This is a modelling study and made no direct use of animals.

### Data Modelling

To test for differences in all sea trout caught at Regional and District scales across all years in the absence of a temporal trend, general linear models (GLMs) with a negative binomial distribution, including year as a repeated measure, were constructed. The test significance is reported as the significance of the deviance explained between the null model including only the repeated measure of “Year” and the model including “Year” as a repeated measure plus “Region” or “District” variables as appropriate. *Post hoc* tests on paired differences were performed using the *emmeans* package (v1.10.3; Lenth 2024) with p-values adjusted using the Tukey method for comparing multiple groups.

Exploratory plots of the data revealed complex, non-linear trends in the sea trout catch timeseries. To describe these trends in the data, variation in the timeseries was decomposed into two components: long-term trends and serial autocorrelation.

The mechanistic processes underlying long-term trends in population sizes are either currently unknown or not understood in enough detail to allow sensible decisions to be made about the shape of the trend line describing temporal change in population size. The modelling approach used to describe long-term trends was therefore entirely driven by the data and a generalised additive modelling (GAM) approach was adopted with a smoothing function identified through the GAM fitting process, as described by Wood (2011). GAM fitting was undertaken using the *mgcv* package (v1.9-1; Wood, 2011) using a quasi-poisson distribution. If the smoothing term from the GAM approach was not significant, a generalised least squares (GLS) model was then used to describe the data, following a square-root transformation of annual sea trout catch. GLS fitting was undertaken using the *nlme* package (v3.1-166; Pinheiro & Bates, 2000; Pinheiro et al., 2024). Pseudo-R^2^ values from the GLS model were calculated as the squared correlation coefficient of the correlation between fitted and observed values from each model.

To account for possible autocorrelation in the time series, autoregressive processes were included in the error term of the models. A smoothing trend and correlated errors can both be fitted to a single time series if a simple structure for the correlation of the error terms is adopted; see Wood (2011, section 7.7.2). Local temporal correlation effects are expected to result from the natural dynamics of the fish population through patterns of reproduction and survival or indeed economic/anthropogenic factors where return rates from one year may have been influenced by the previous year(s). Autoregressive processes provide a simple means of modelling this, protecting the estimation and interpretation of the long-term trend from the effects of local temporal correlation. Identification of the optimal autoregressive process for describing serial correlation across all spatial scales was undertaken through comparison (likelihood ratio testing and AIC comparison) of competing models (AR1 vs AR2 vs AR0, respectively models comprising a one year temporal autocorrelation offset, a two year offset and a naïve model assuming no temporal autocorrelation). The optimal AR process was identified as the AR2, which best captured the serial correlation in all the times series at the National, Regional and District scales. This provides a more flexible description of correlation across the series than the simple, rapidly diminishing geometric decay of AR1. As the return times for sea trout to the freshwater environment in Scotland can be anything from around one to in excess of six years from hatching, the greater adaptability of the AR2 autocorrelation structure is helpful in capturing the serial correlation while attributing the remaining variation to longer term trends.

To identify periods of statistically significant change across the GAM trendline, we used the method of finite differences to estimate the rate of change (slope/derivative) in the fitted smoother(adapted code from Simpson, 2014). The fitted smoothing function is evaluated at a number of equidistant points along the time axis. The derivative of the function can then be estimated from the differences of the function estimates at adjacent time points divided by the length of the time axis intervals. The standard errors of the original smoothing curve can also be propagated through this calculation to provide standard errors for the derivative. This allows sections of the smoothing estimate to be identified where the derivative is significantly different from zero. This approach was replicated over the three spatial scales (Wood et al., 2017; Pedersen et al 2019). The modelling approach accounts for serial autocorrelation through an AR error term. This attributes short term patterns in plots of the raw data to serial correlation rather than systematic changes in the mean. All analysis was undertaken in R v4.4.1 (R Core Team, 2024).

A sensitivity analysis investigating the effects of excluding the data from 2020 (when restrictions imposed during the COVID pandemic had the potential to disproportionately impact on sea trout catches) showed no material effect on the relationships described when including 2020 (Supplementary Information Figure S1). In addition, the proportion of angling returns made for 2020 (0.927) was broadly in line with preceding (e.g. 2017=0.945, 2018=0.932, 2019=0.940) and subsequent years (2021=0.938, 2022=0.936, 2023=0.914) (Scottish Government, Marine Directorate, 2024). Data from 2020 were therefore included in all analyses.

## RESULTS

### National Scale Sea trout catches

At the national scale, annual sea trout catch varied from a maximum of 66,544 in 1966 (second highest year was1965, where catch was 62,865, third highest 1964, catch=55,152) to a minimum of 16,696 in 2021 (second lowest year was 2020 catch=17,504, third lowest 2023 catch=17,930). The mean number of fish caught across all rivers and all years was 34,463 (95% CI=31966.6, 36909.3) with a coefficient of variation of 30.5.

The annual sea trout catch from across Scotland showed a significant decrease across the 72-year study period (Figure 2). The model explained 60.3% (adjusted R^2^=0.603) of the variation in annual sea trout catch. The smoothed term for Year was strongly statistically significant (EDF=2.5, F=21.8, P<0.0001) indicating a non-linear trend which changed from a model predicted maximum in 1952 of 48,405 (SE±4,086) to a minimum in 2023 of 22,871 (SE±2,365). This represents a reduction of 52.8% in reported annual sea trout catch over the study period (from 1952).

**Figure 2.**
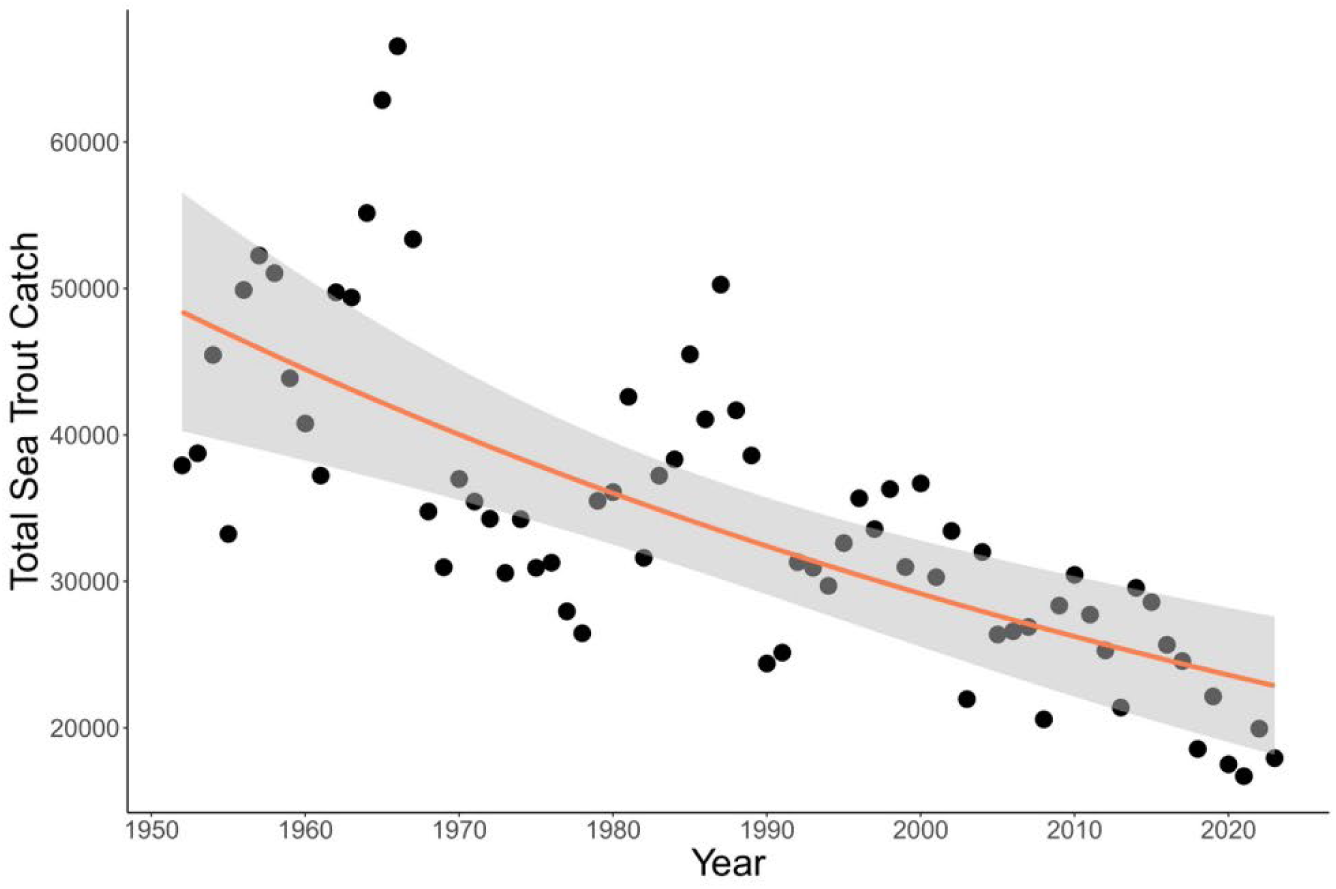
Recorded total annual rod catches of sea trout from across Scotland (the National scale) from the 64 Districts included in the data set. Fitted line from GAM model in orange with the shading indicating the 95% confidence interval.

### Regional Scale Sea Trout catches

#### Catch differences between Regions, all years combined

There were significant differences in total annual sea trout catch between Regions (χ^2^_(8)_ =577.14, p<0.0001) and statistically significant differences detected in pairwise comparisons between Regions (Figure 3).

**Figure 3.**
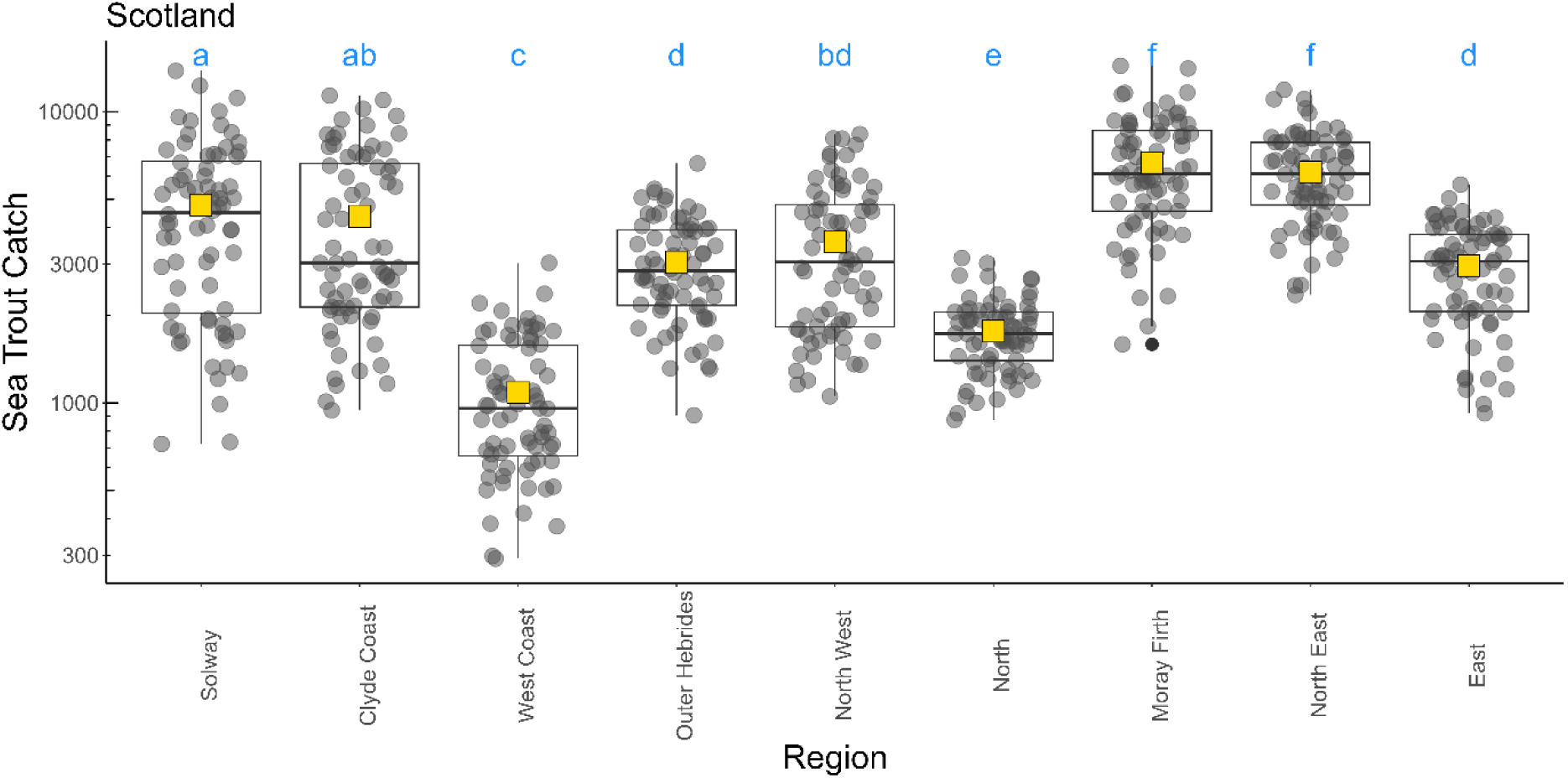
Mean (yellow square), median, interquartile range, minimum and maximum annual sea trout catches across all years between Regions (catch plotted on the log10 scale). Shared alpha character indicates no statistical difference (p>0.05 (after correcting for multiple pairwise comparisons)).

Overall, mean sea trout catches across all years, ranged from a maximum of 6,655 (in the Moray Firth Region) to a minimum of 1,089 (in the West Coast Region) (Table 1). The coefficient of variation ranged from 27.9% (in the North Region) to 64.6% (in the Clyde Coast Region) (Table 1), indicating considerable differences in the variability of sea trout catches between different Regions with the Clyde Coast Region showing the highest variability in sea trout catches over the study period.

**Table 1.** Mean reported sea trout annual rod catch (number, with standard error in parenthesis), coefficient of variation, lower and upper 95% confidence limits, minimum and maximum total annual catch and the years in which the minimum and maximum occurred in each of the nine Regions in Scotland over the period of 1952 to 2023.

| Region | Mean (SE) | CV | Lower | Upper | Min Catch | Min Year(s) | Max Catch | Max Year |
| --- | --- | --- | --- | --- | --- | --- | --- | --- |
| Solway | 4780.1 (345.4) | 61.3 | 4091 | 5469 | 723 | 2021 | 13,819 | 1966 |
| Clyde Coast | 4377.7 (333.3) | 64.6 | 3713 | 5042 | 944 | 2006 | 11,361 | 1965 |
| West Coast | 1089 (67.1) | 52.3 | 955 | 1223 | 292 | 1991 | 3,026 | 1964 |
| Outer Hebrides | 3049.4 (139.6) | 38.8 | 2771 | 3328 | 907 | 1980 | 6,646 | 2015 |
| North West | 3572.2 (235) | 55.8 | 3104 | 4041 | 1,053 | 2003 | 8,377 | 1965 |
| North | 1763.1 (58.1) | 27.9 | 1647 | 1879 | 875 | 1952 | 3,148 | 2011 |
| Moray Firth | 6655.2 (328.6) | 41.9 | 6000 | 7310 | 1,589 | 2020 | 14,386 | 1965 |
| North East | 6197.9 (252) | 34.5 | 5696 | 6700 | 2,350 | 1961 | 11,888 | 1957 |
| East | 2951.9 (129.5) | 37.2 | 2694 | 3210 | 921 | 1960 | 5,623 | 2011 |

#### Patterns of temporal change

The pattern of change in sea trout catches show considerable inter-regional variation which, for six of the nine Regions, differed from the trend shown at the national scale (Figure 4). For the three Regions, Clyde Coast (model R^2^_adj_=0.565; smoothing term EDF=1, F=13.8, p=0.0004), West Coast (model R^2^_adj_=0.577; smoothing term EDF=1, F=31.9, P<0.0001) and North West, (model R^2^_adj_=0.769; smoothing term EDF=1, F=95.6, P<0.0001) there was a significant decline in sea trout catches across the timeseries, reflecting the national trend. Modelled estimates of change (Table S3) for each of these Regions represented a total reduction since the start of the data series in 1952 of 78.2% for the Clyde Coast (modelled catch (mc) in 1952 = 8,405 (SE±1,596) to mc=1,832 (SE±534) in 2023), 73.4% for the West Coast (mc=1,939 (SE±217) in 1952 to mc=515 (SE±83) in 2023) and 81.4% for the North West (1952 mc=7,340 (SE±571) to mc=1,368 (SE±165) in 2023), equating to an annual loss rate of 1.09%, 1.02% and 1.13% respectively.

**Figure 4.**
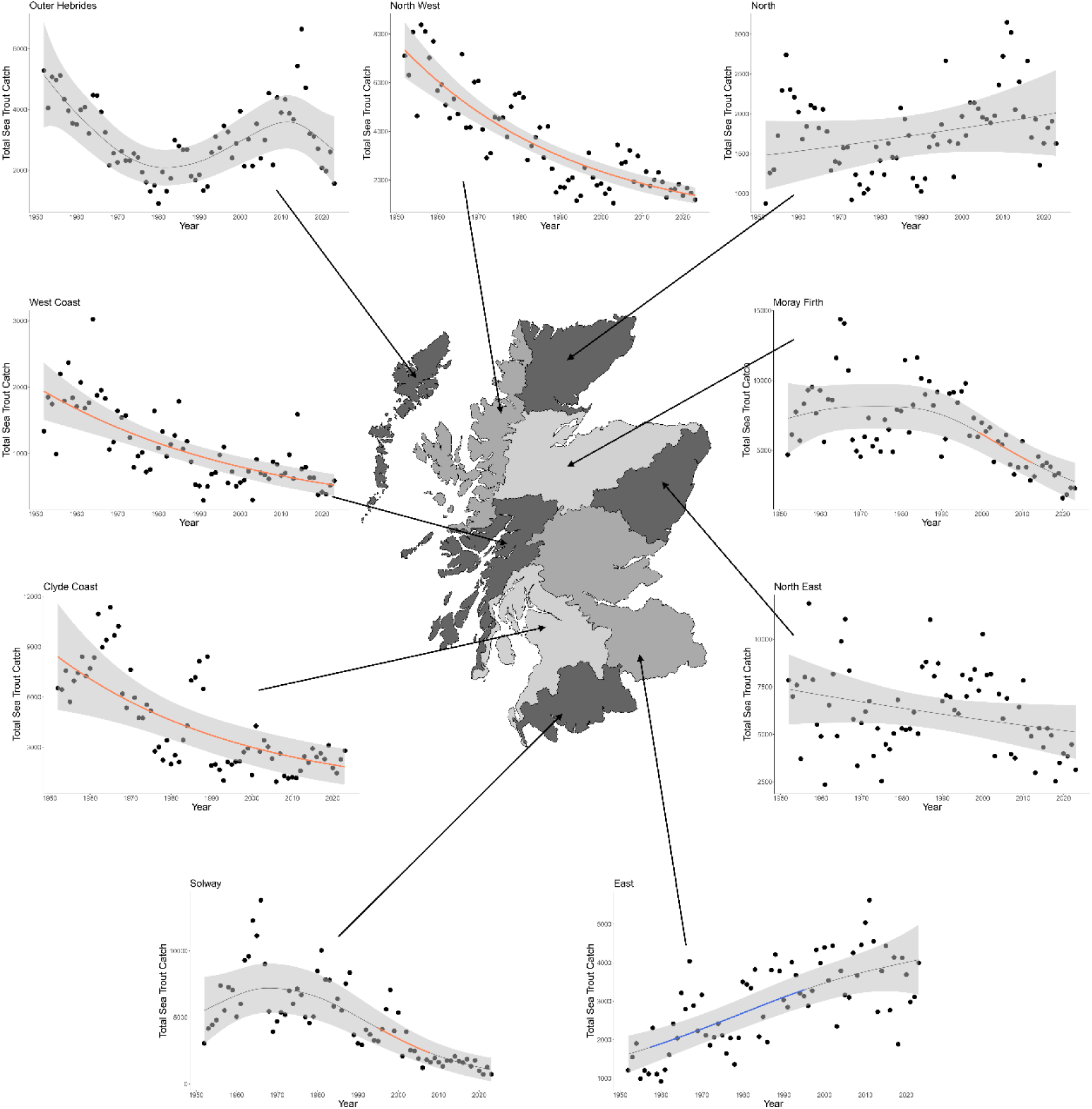
Annual recorded rod catches of sea trout between 1952 and 2023 from the 64 Districts included in the data set at the Regional scale (clockwise from bottom left: Solway, Clyde Coast, West Coast, Outer Hebrides, North West, North, Moray Firth, North East, and East). Solid lines represent the predicted relationship (including the 95% confidence interval as shaded area) from the model (Table 1). Periods of statistically significant increase (blue) and decrease (orange) in rod catches are highlighted. Map redrawn from Scottish Government (2025).

For two Regions, the Solway (model R^2^_adj_=0.603; smoothing term EDF=2.5, F=9.17, p=0.0001) and the Moray Firth (model R^2^_adj_=0.460; smoothing term EDF=2.4, F=7.73, p=0.0008), there were significant declines detected for part of the time series, Solway from 1994 (mc=4,229 SE±660) to 2007 (mc=2,262 SE±479) and Moray Firth from 1999 (mc=6,308 SE±685) to 2013 (mc=4,082 SE±590) (Figure 4, Table S3) representing a loss in reported sea trout catches of 46.5% and 35.3% respectively over these periods. One Region, East (model R^2^_adj_=0.491; smoothing term EDF=1.6, F=18.7, p<0.0001), showed a statistically significant increase in catches between 1957 (mc=1,804 SE±219) to 1991 (mc=3,302 SE±221) representing an increase in sea trout catches of 45.4%, equating to an annual gain rate of 1.3%. across this period. The remaining three Regions, the Outer Hebrides (model R^2^_adj_=0.546; smoothing term EDF=4.0, F=5.08, P=0.002), the North (model R^2^_adj_=0.077; smoothing term EDF=1, F=1.65, P=0.203) and the North East (model R^2^_adj_=0.057; smoothing term EDF=1, F=2.60, P=0.111), showed no statistically significant periods of change across the timeseries. For two Regions, the North and the North East, the smoothing term in the GAM was not significant. GLS models for each Region also showed a non-significant linear trend (North pseudo-R^2^=0.087, t=1.15_(72,70)_, p=0.254; North East pseudo-R^2^=0.076, t=1.49_(72,70)_, p=0.142).

Based on the adjusted R^2^ statistic from the model, the amount of variation explained by temporal change (i.e. variation explained by serial autocorrelation plus the GAM smoother) ranged from very low (R^2^ = 0.077 for the North Region) to explaining almost 80% of the variation in sea trout catches (R^2^ =0.767 for the North West Region; Table 2).

**Table 2.**
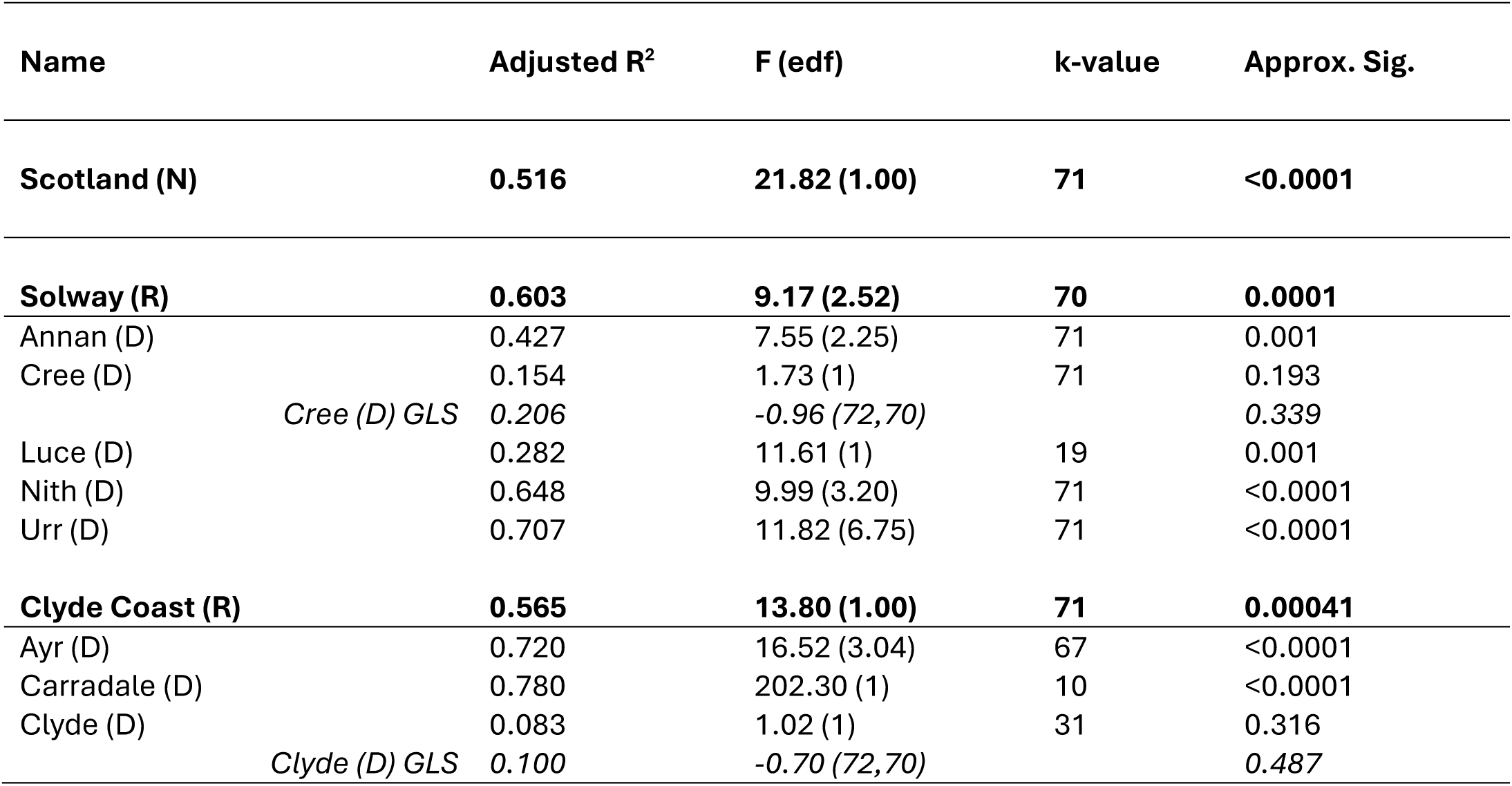
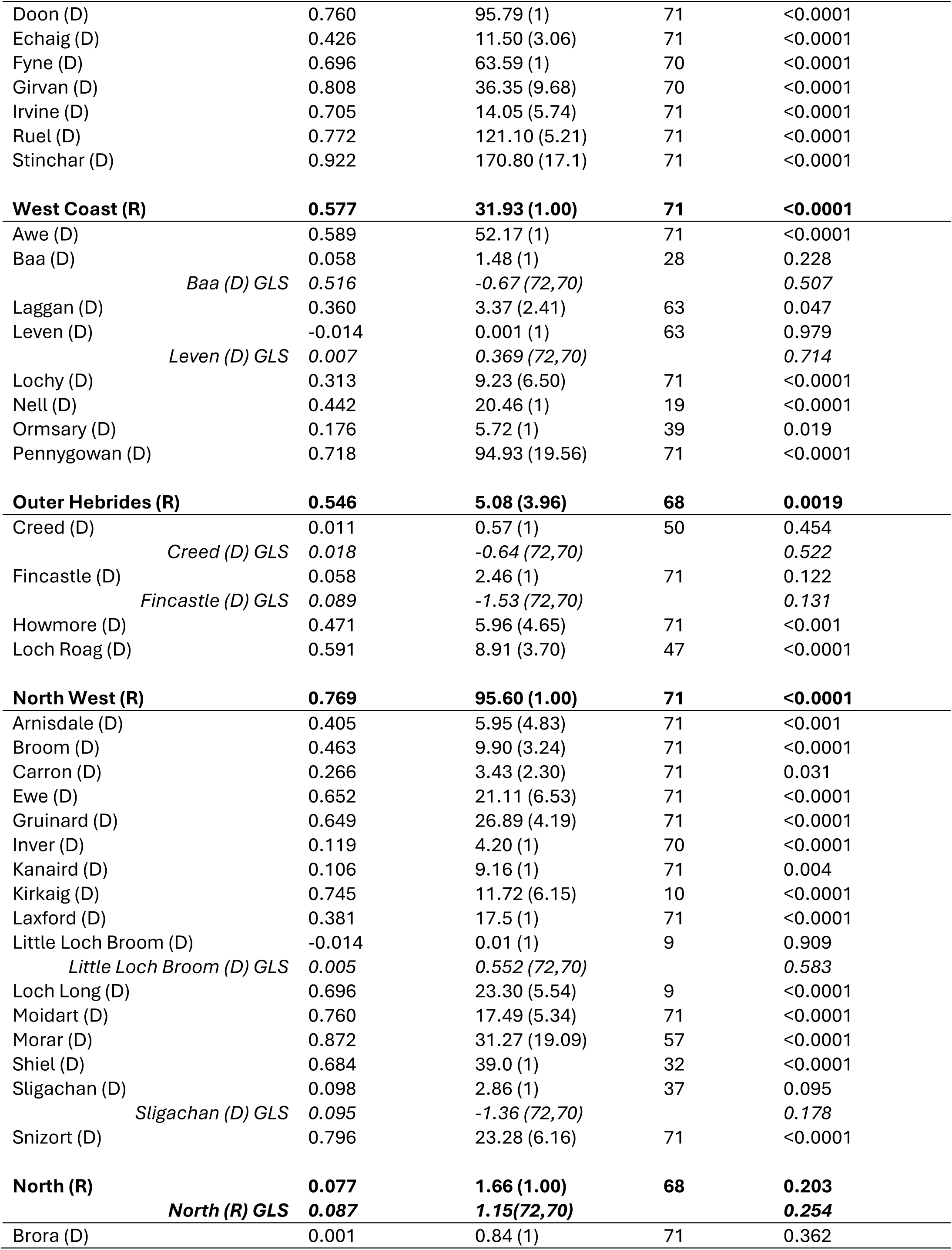
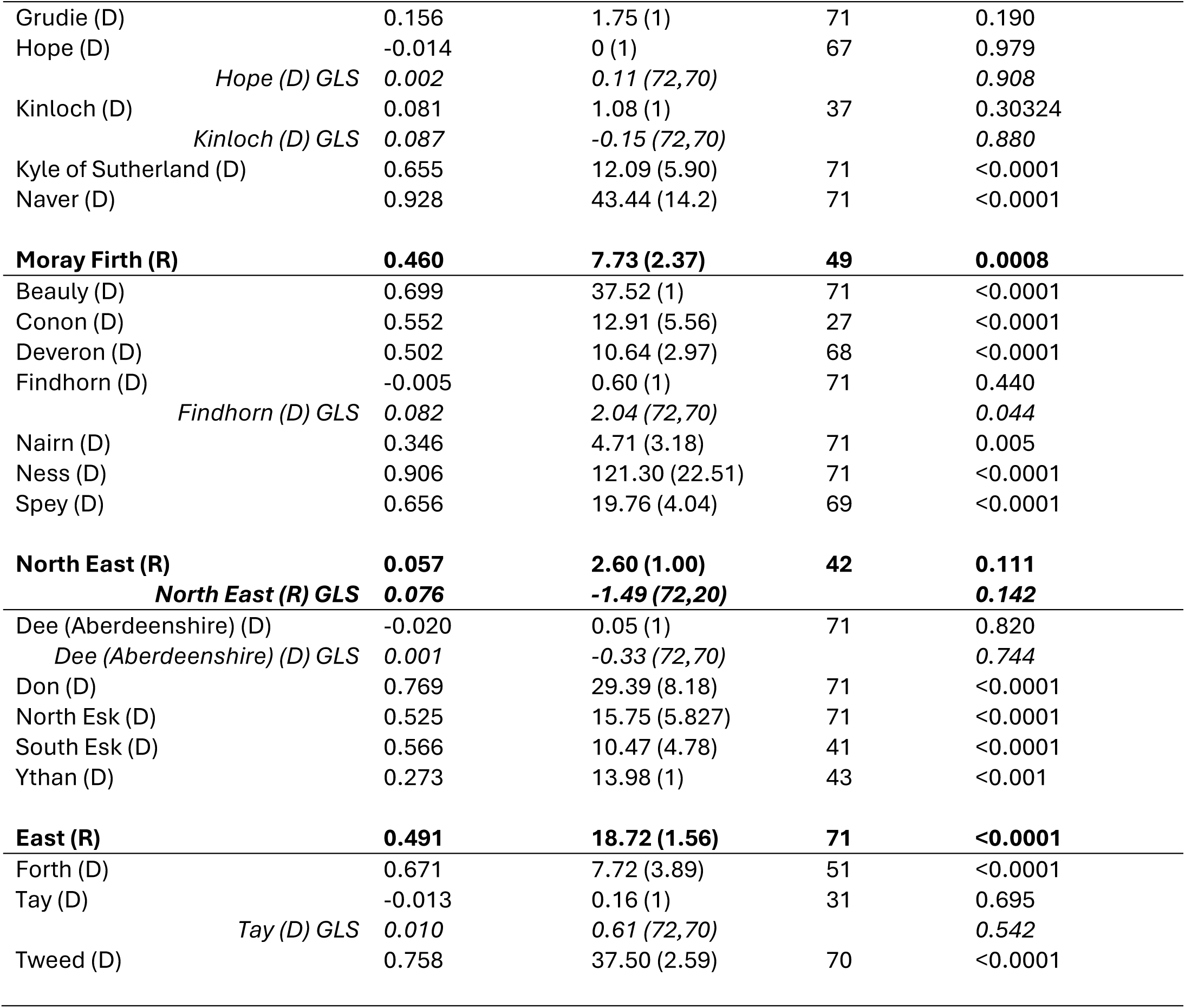
Summary results from the statistical testing. Model adjusted R^2^, the F-value and associated effective degrees of freedom (edf), the number of basis function for the GAM (k-value) and the approximate significance of the combined smoothing term and the chosen autoregressive process, for each District (D) and Region (R) in this study and Nationally (N). For those tests where the smoother was not significant, the results from the GLS test have been included in italics detailing pseudo-R^2^, test statistic (t) and degrees of freedom and associated level of significance.

Estimation of effects at national, regional and district levels are inevitably affected by the amount of information available at the different scales being considered. The sample sizes contributing to the estimation of each temporal trend decrease markedly from national to regional to district levels. This is generally sufficient to identify the presence of trend and its broad characteristics but the identification of more detailed features may be more challenging. This problem is particularly acute when identifying periods of time during which trends are increasing or decreasing, as these local calculations further reduce the effective sample size. The periods identified in this study are therefore restricted to those where the local trend is particularly strong.

#### Sea trout catches at the District scale

Each of the nine Regions comprise a number of Districts ranging from three for the East Region to 16 for the North West Region (Figure 1; Table S1). Across all Districts, statistical testing identified 57 significant periods of marked change in 41 Districts (23 Districts did not show any significant change). Of these 40 were sea trout catch declines and 17 were catch increases (Table 2). We examine the patterns of change in sea trout catches at the District scale that comprise each of the nine Regions.

#### Districts in the Solway Region

The Solway Region (the most south-westerly of the nine Regions; Figure 1a) comprises eight reporting Districts, five of which were included in the cleaned dataset (Table S1). There were significant differences in total annual sea trout catch between different Districts within the Solway Region (χ^2^_(4)_=586.0, p<0.0001) and pairwise comparisons showed that each District was statistically different from the others at p=0.05 level (Figures 5a,6). Within the Solway Region, the District with the highest mean annual catch was the Nith (2,812) and the lowest was the Urr (167) (Table S2).

**Figure 5.**
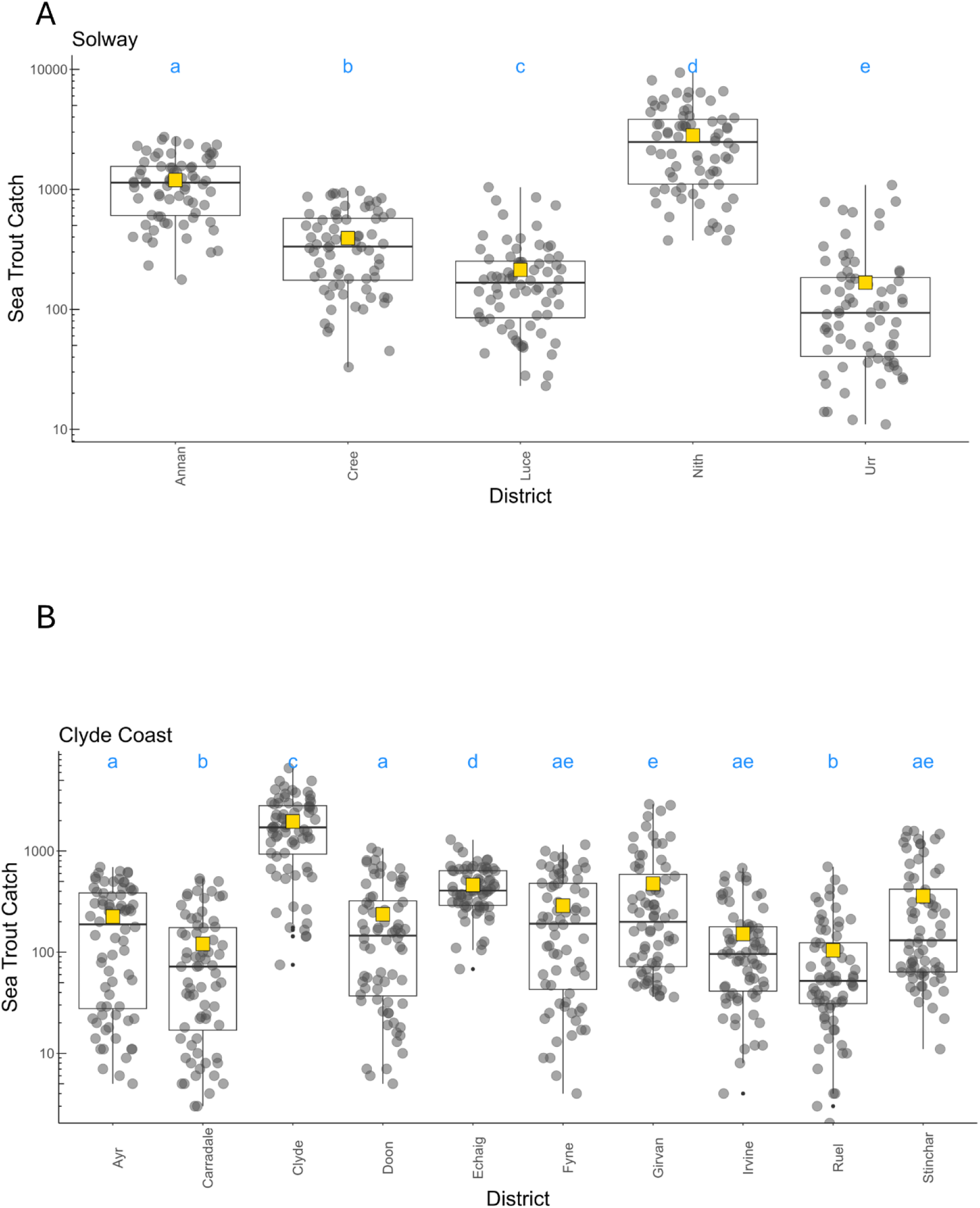
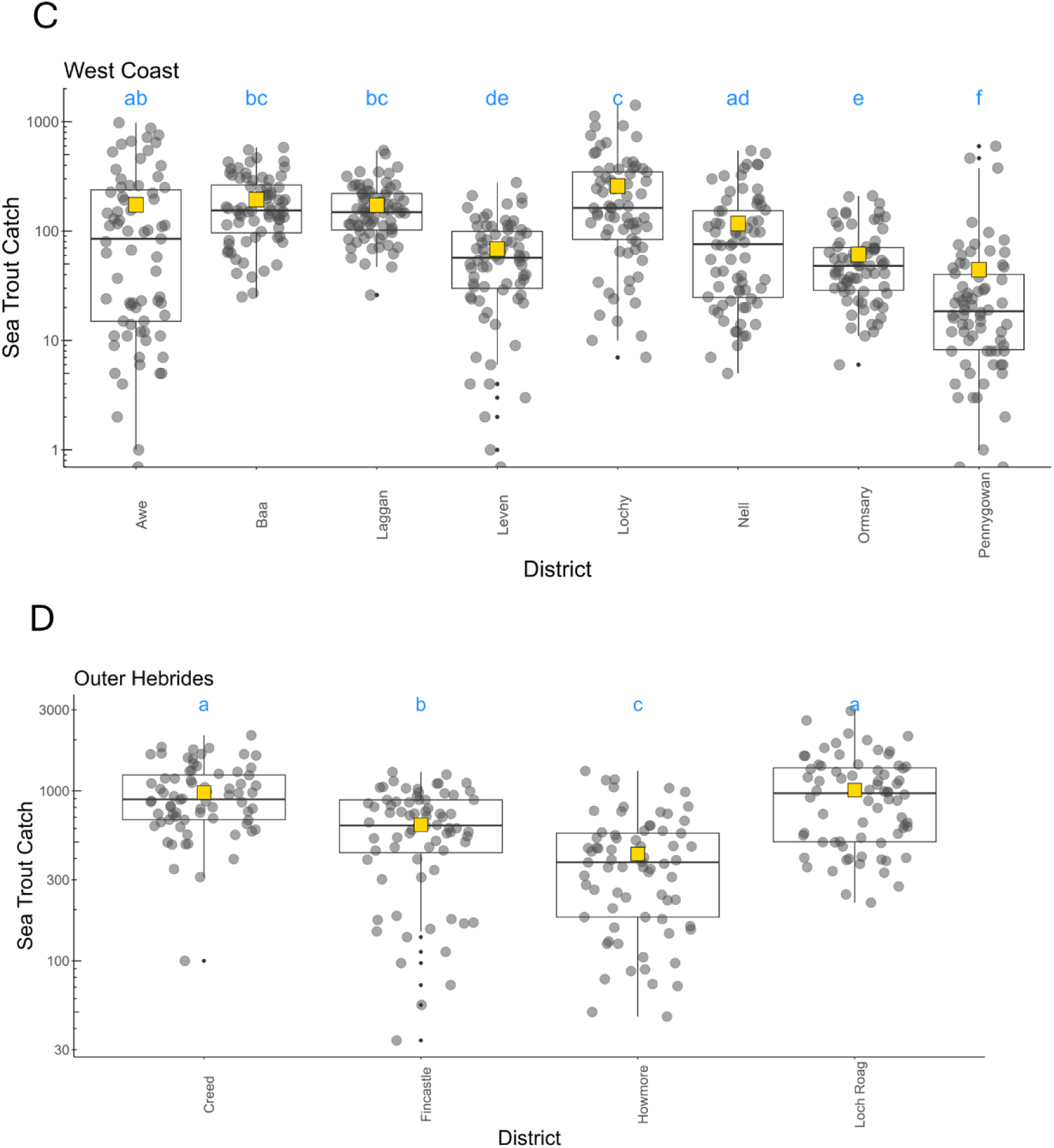
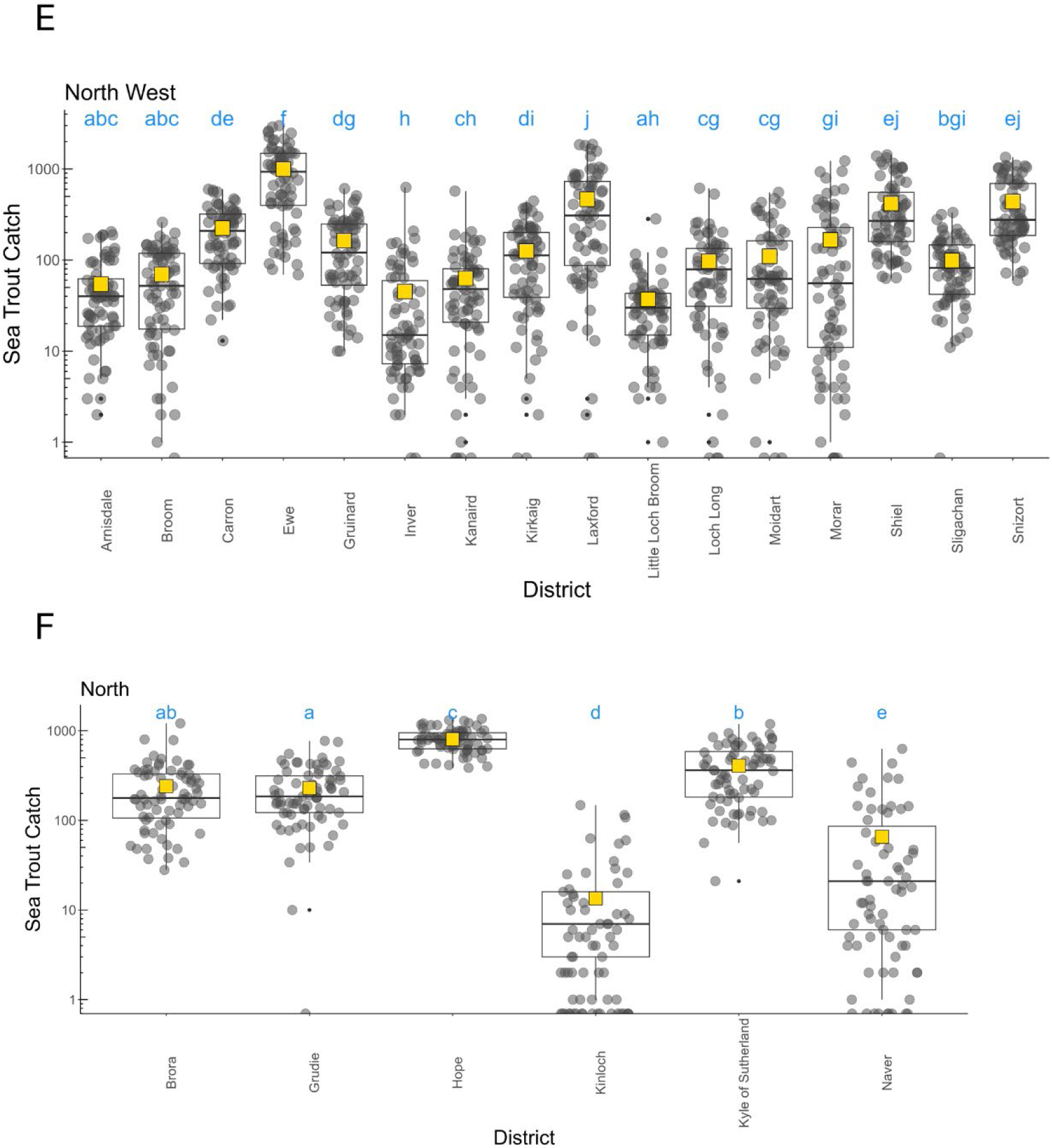
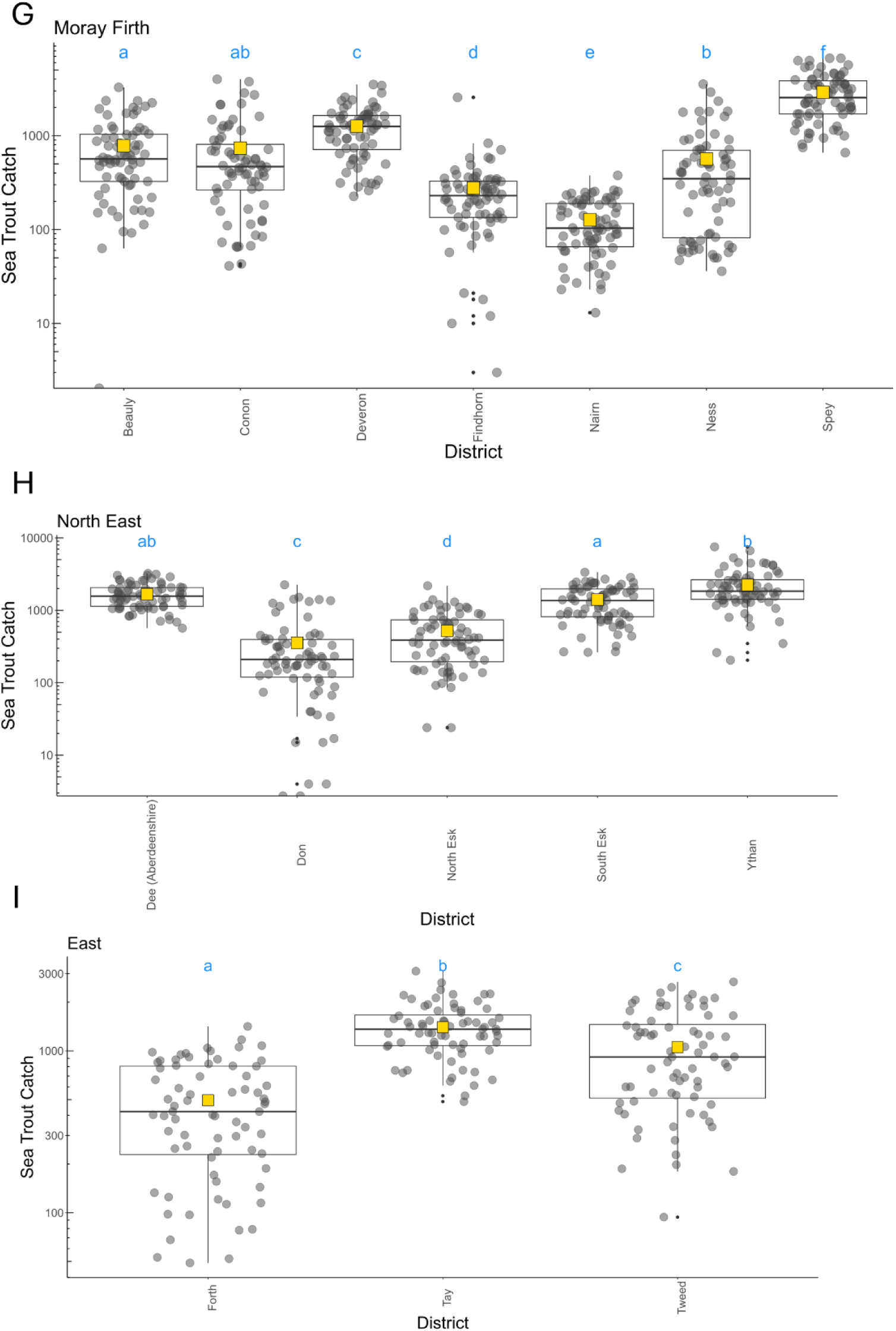
Mean (yellow square), median, interquartile range, minimum and maximum annual sea trout catch (plotted on the log10 scale) across all years for the Districts within the nine Regions: a = Solway, b = Clyde Coast, c = West Coast, d = Outer Hebrides, e = North West, f = North, g = Moray Firth, h = North East, and i = East. Total annual catch is plotted on). Shared alpha character indicates no statistical difference (p>0.05 (after correcting for multiple pairwise comparisons)).

**Figure 6.**
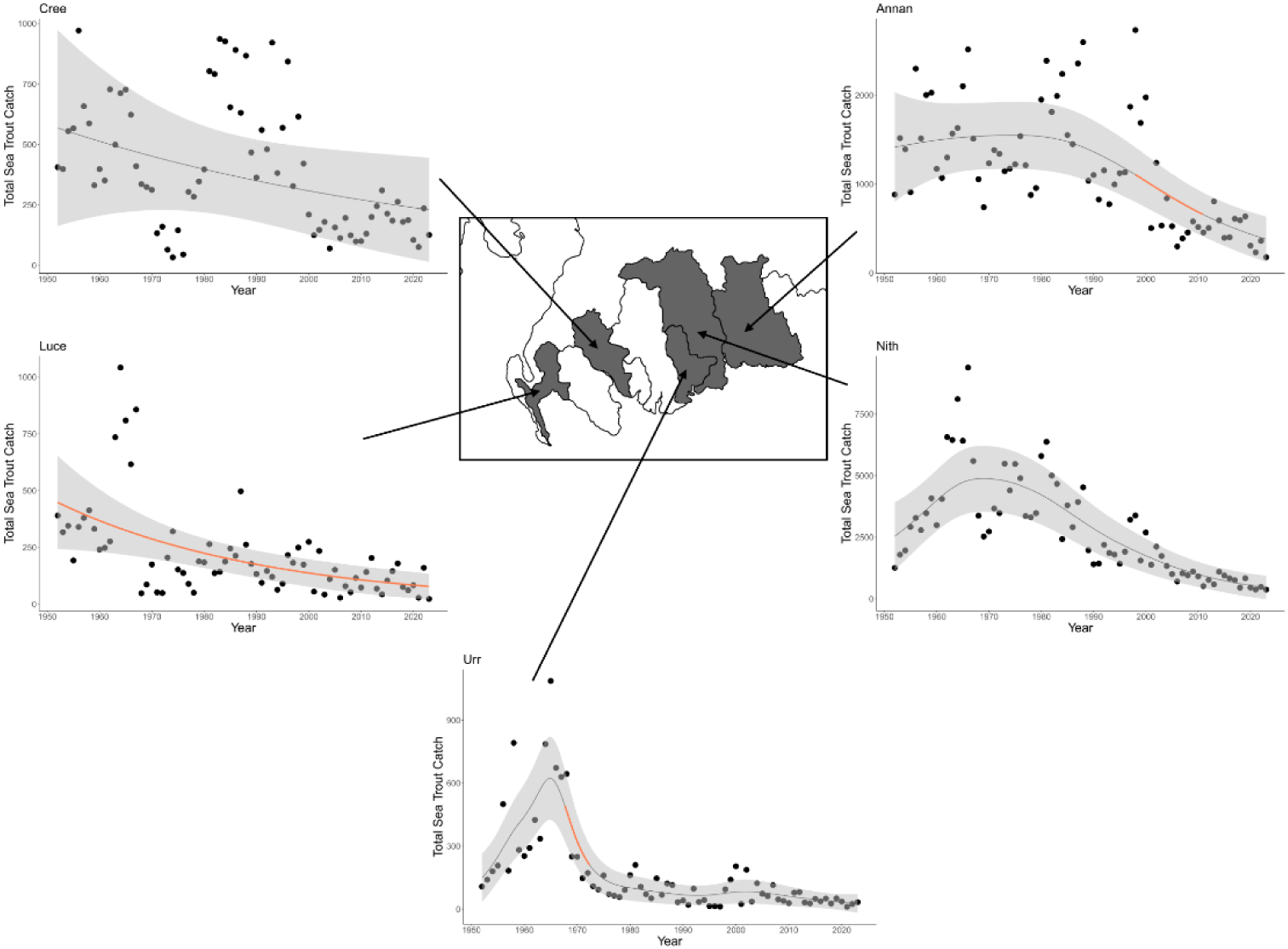
Rod catches of sea trout from each of five Districts in the Solway regions (from east to west: the Annan, Nith, Urr, Cree and Luce Districts) between 1952 and 2023. Solid fitted lines are the model predicted values, dotted lines are 95% confidence limits around these estimates. Solid lines represent the predicted relationship (including the 95% confidence interval as shaded area) from the model (Table S4). Periods of statistically significant increase (blue) and decrease (orange) in rod catches are highlighted. Map redrawn from Scottish Government (2024b).

Three Districts, Anna, Luce, and Urr, showed statistically significant periods of decline in sea trout catches (Figure 6, Table 2, Table S4). Significant decline across the whole timeseries was identified in the Luce District, in the Annan from 1997 to 2012, and in the Urr from 1966 to 1971. There was no statistical change in catches over the timeseries detected in the Cree or Urr Districts (Figure 6, Table 2). Of the five Districts within the Solway Region, the Annan District was the only District showing a similar pattern of change to that of the Solway Region (ie a significant decline from 1993 to 2008).

#### Districts in the Clyde Coast Region

The Clyde Coast Region comprises 12 reporting Districts, ten of which were included in the cleaned dataset (Figure 7). There were significant differences in total annual sea trout catch between different Districts within the Clyde Coast Region (χ^2^_(9)_ =687.7, p<0.0001). Statistical difference was identified between 31 of the pairwise comparisons of Districts (Figure 5b). Within the Clyde Coast Region, the District with the highest mean annual catch was the Clyde (1,958) and the lowest was the Ruel (104) (Table S2).

**Figure 7.**
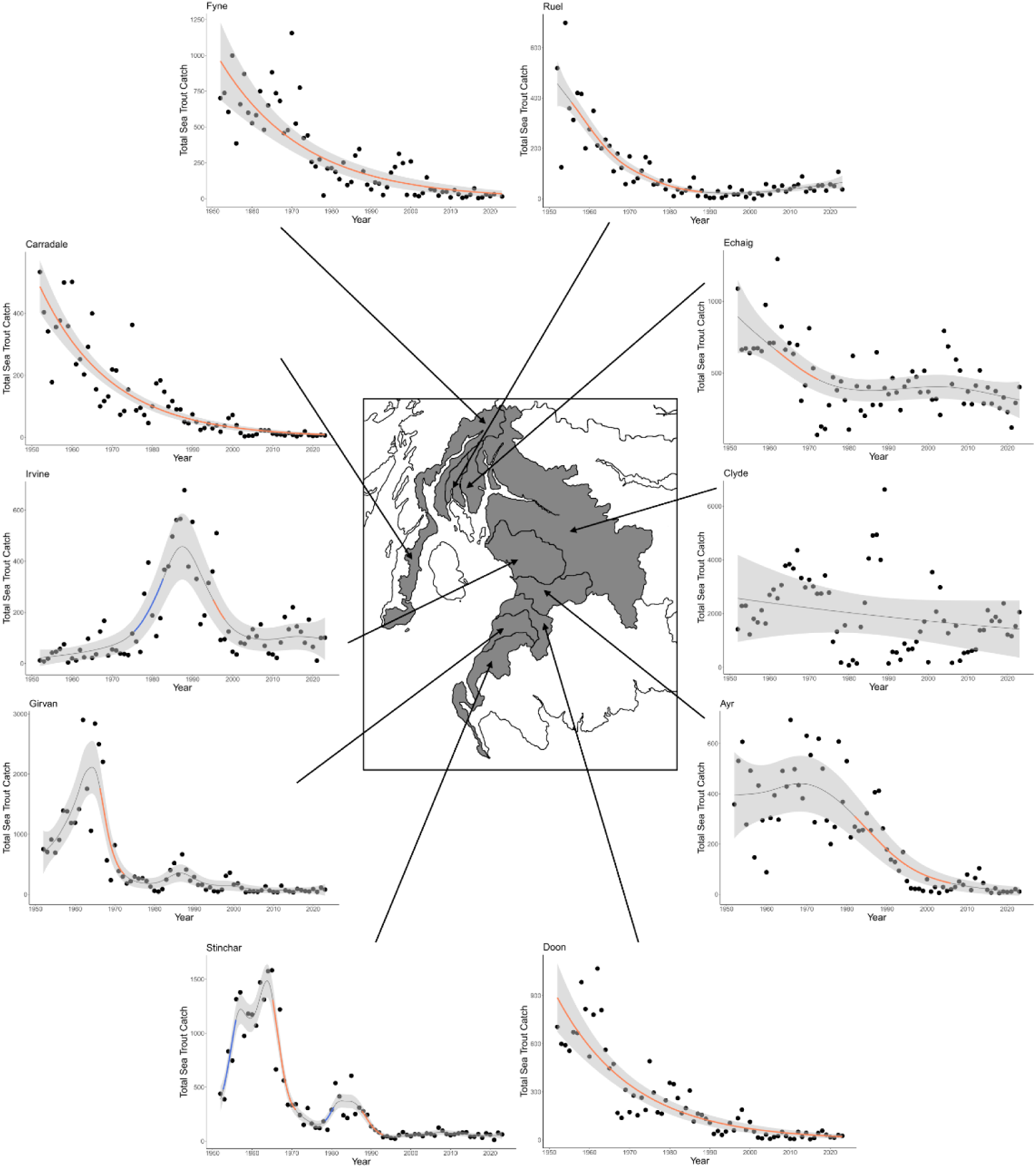
Rod catches of sea trout from each of 10 catchments in the Clyde Coast Region (from south to north: the Stinchar, Girvan, Doon, Ayr, Irvine, Clyde, Carradale, Eachaig, Ruel & Fyne) between 1952 and 2023. Solid lines represent the predicted relationship (including the 95% confidence interval as shaded area) from the model (Table S4). Periods of statistically significant increase (blue) and decrease (orange) in rod catches are highlighted. Map redrawn from Scottish Government (2025).

Thirteen statistically significant periods of marked change were identified in nine Districts (Figure 7, Table 2, Table S4). Decline in catches were identified in nine Districts (Figure 7); the Ayr (1982 to 2004), the Carradale (across the whole timeseries), the Doon (across the whole timeseries), the Echaig (1959 to 1973), the Fyne (across the whole timeseries), the Girvan (1966 to 1972), the Irvine (1996 to 1997), the Ruel (1955 to 1987) and the Stinchar (1965 to 1971 and 1987 to 1992). There was a period of significant catch increase in the Irvine (1976 to 1981). Two periods of significant catch increase were also identified in the Stinchar (1952 to 1955 and 1979 to 1980). There was no statistical change in catches over the timeseries detected in the Clyde District (Figure7, Table 2, Table S4). Of the ten Districts within the Clyde Coast Region, three Districts, Carradale, Doon, and Fyne showed patterns of change similar to this identified at the Regional scale (i.e. significant decline in sea trout across the whole timeseries).

#### Districts in the West Coast Region

The West Coast Region comprises 16 reporting Districts, eight of which were included in the cleaned dataset (Figure 8). There were significant differences in total annual sea trout catch between different Districts within the West Coast Region (χ^2^_(9)_ =212.7, p<0.0001). Statistical difference was identified in 20 of the pairwise comparisons of Districts (Figure 5c). Within the West Coast Region, the District with the highest mean annual catch was the Lochy (258) and the lowest was the Pennygowan (44) (Table S2, Table S4).

**Figure 8.**
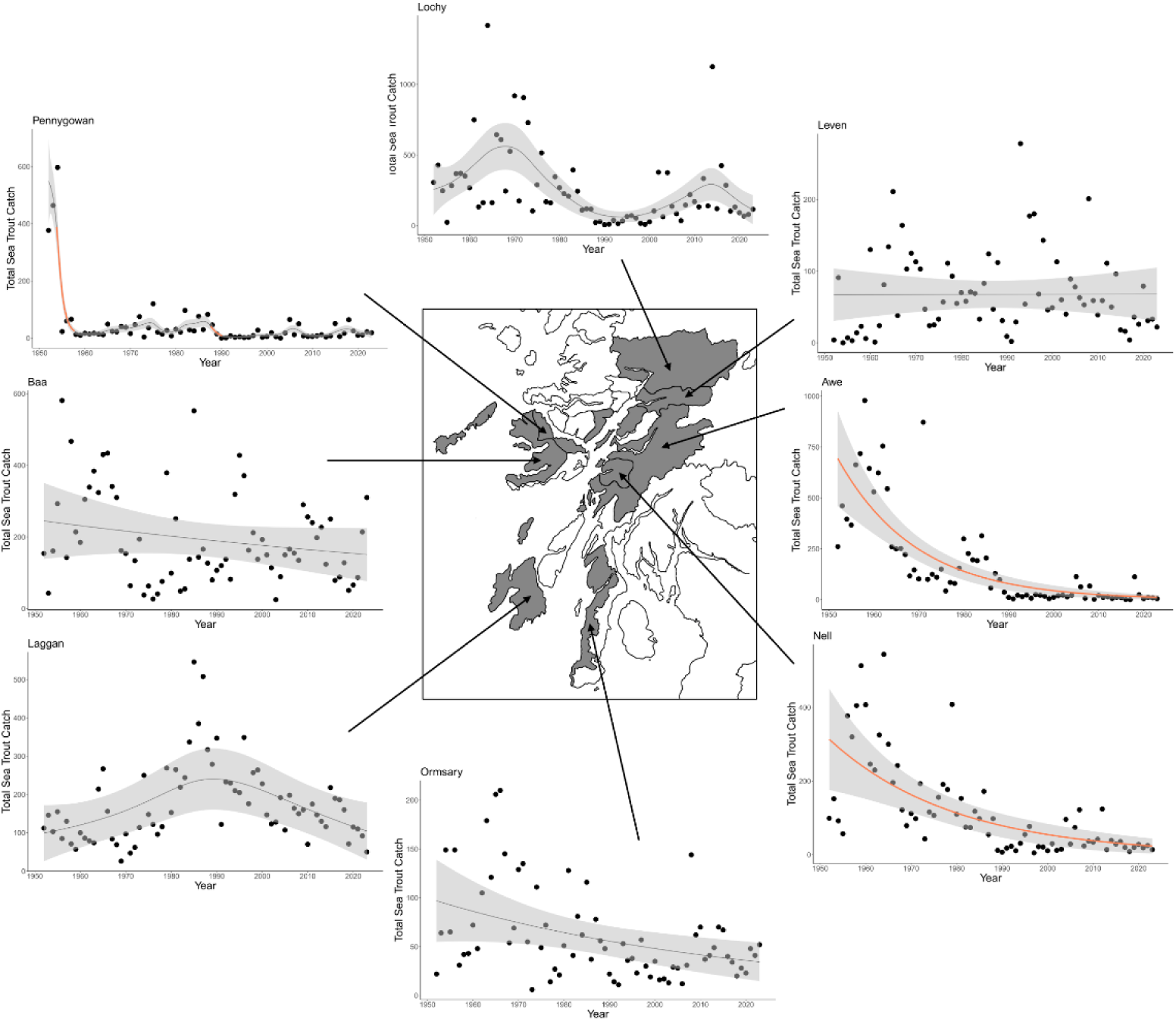
Rod and line catches of sea trout from each of the eight Districts in the West Region (from south to north: the Ormsary, Laggan, Nell, Awe, Baa, Pennygowan, Leven and Lochy) between 1952 and 2023. Solid lines represent the predicted relationship (including the 95% confidence interval as shaded area) from the model (Table S4). Periods of statistically significant increase (blue) and decrease (orange) in rod catches are highlighted. Map redrawn from Scottish Government (2025).

Four statistically significant periods of marked decline were identified in three Districts (Figure 8, Table 2, Table S2, Table S4). These were in the Awe (across the whole timeseries), the Nell (across the whole timeseries), and the Pennygown (1953 to 1957 and 1987 to 1990) (Figure 8). Five Districts, Baa, Laggan, Leven, Lochy, and Ormsary showed no periods of statistically significant change (Figure 8). Of the eight Districts within the West Coast Region, two Districts, Awe and Nell showed similar patterns of change identified at the Regional scale (i.e. significant decline in sea trout across the whole timeseries).

#### Districts in the Outer Hebrides Region

The Outer Hebrides Region comprises eight reporting Districts, four of which were included in the cleaned dataset (Figure 9). There were significant differences in total annual sea trout catch between Districts within the Region (χ^2^_(3)_ =94.77, p<0.0001) Statistical difference was identified in five of the pairwise comparisons of Districts (Figure 5d). Within the Outer Hebrides Region, the District with the highest mean annual catch was the Loch Roag (1,011) and the lowest was the Howmore (425) (Table S2, Table S4).

**Figure 9.**
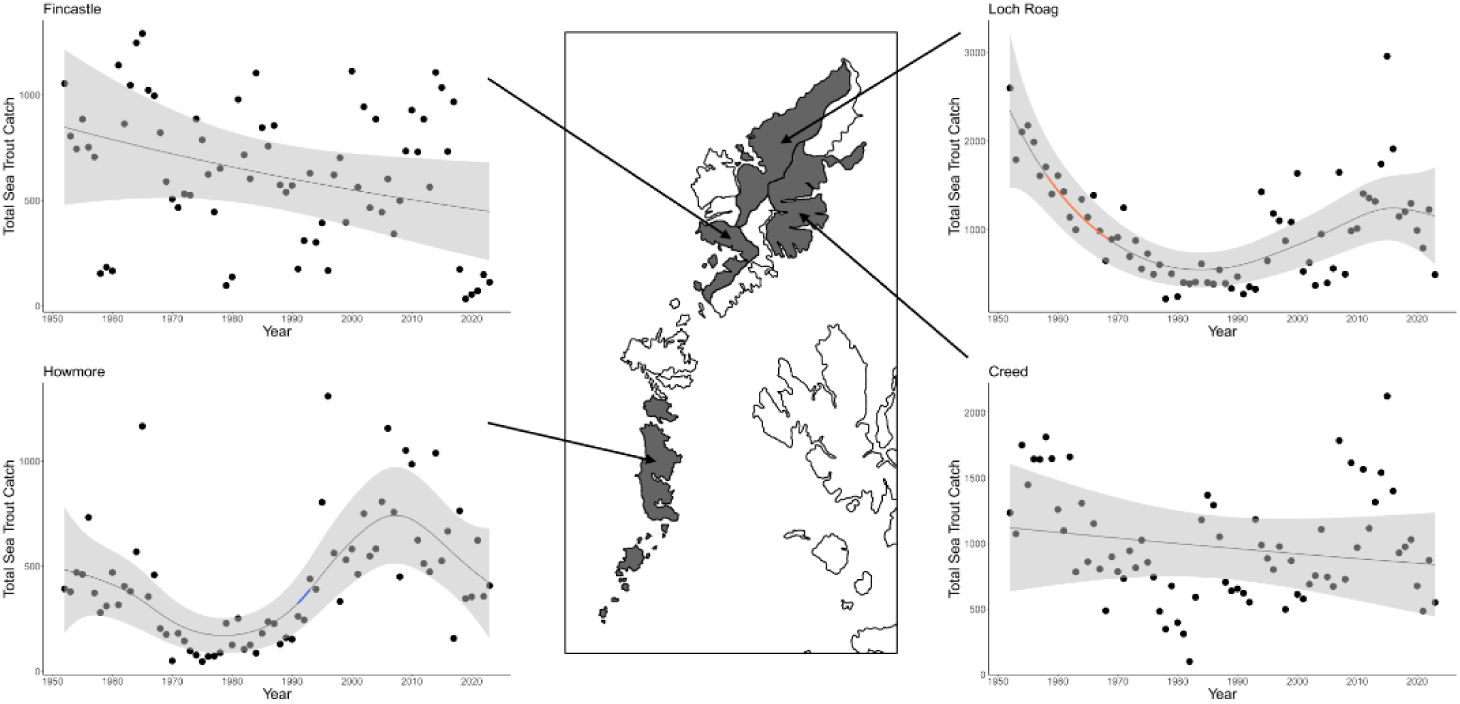
Rod catches of sea trout from each of the four Districts in the Outer Hebrides Region (from south to north: the Howmore, Fincastle, Creed and Loch Roag Catchments) between 1952 and 2023. Solid lines represent the predicted relationship (including the 95% confidence interval as shaded area) from the model (Table S4). Periods of statistically significant increase (blue) and decrease (orange) in rod catches are highlighted. Map redrawn from Scottish Government (2025).

Two statistically significant periods of marked change were identified in two Districts (Figure 9, Table 2, Table S2, Table S4). Decline in catches were identified in the Loch Roag District (1958 to 1968) and an increase in catches identified in the Howmore (1990 to 1993) (Figure 9). There was no statistical change in catches over the time series detected in the Creed or the Fincastle (Figure 9, Table 2). Of the four Districts within the Outer Hebrides Region, two, the Creed and the Fincastle, showed similar patterns of change identified at the Regional scale (i.e. no significant change across the timeseries).

#### Districts in the North West Region

The North West Region comprises 28 reporting Districts, 16 of which were included in the cleaned dataset (Figure 10). There were significant differences in total annual sea trout catch between different Districts within the Region (χ^2^_(15)_ =833.59, p<0.0001).

**Figure 10.**
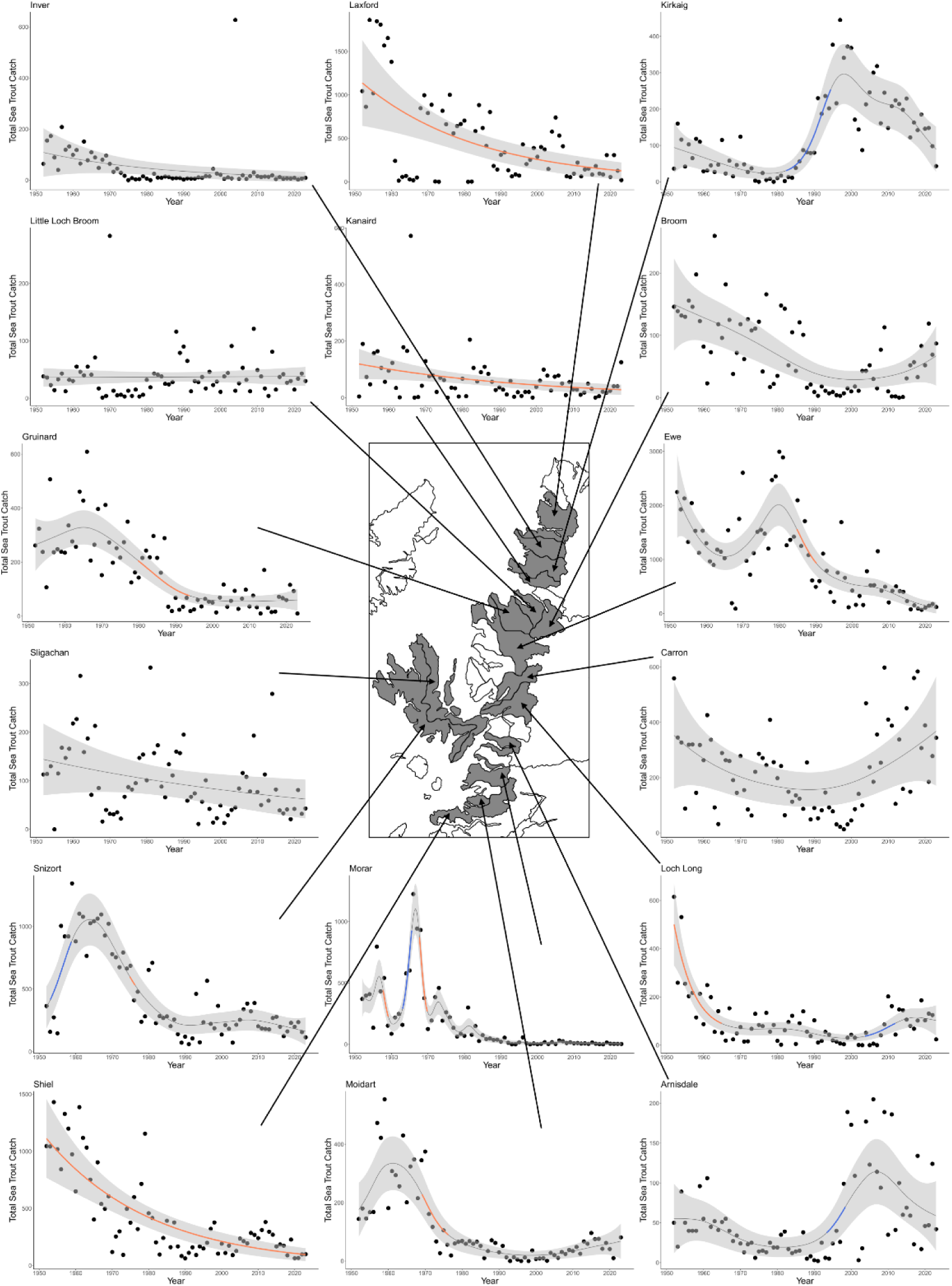
Rod catches of sea trout from each of the 16 Districts in the North West Region (from south to north: the Sheil, Moidart, Morar, Arnisdale, Snizort, Loch Long, Sligachan, Carron, Ewe, Broom, Gruinard, Little Loch Broom, Kanaird, Kirkaig, Inver & Laxford) between 1952 and 2023. Solid lines represent the predicted relationship (including the 95% confidence interval as shaded area) from the model (Table S4).

Statistical difference was identified between 86 pairwise comparisons of Districts (Figure 5e). Within the North West Region, the District with the highest mean annual catch was the Ewe (996) and the lowest was the Little Loch Broom (37) (Table S2, Table S4).

Fifteen statistically significant periods of marked change were identified in eleven Districts (Figure 10, Table 2, Table S2, Table S4). Decline in catches were identified in eight Districts (Figure 10), the Ewe (1984 to 1990), the Gruinard (1979 to 1993), the Kanaird (across the whole timeseries), the Laxford (across the whole timeseries), Loch Long (1952 to 1964), the Moidart (1969 to 1976), the Morar (1957 to 1959 and 1967 to 1969), the Shiel (across the whole timeseries), and the Snizort (1974 to 1976). The Morar and the Snizort also showed periods of significant increase (1963 to 1965 and 1953 to 1958, respectively). Significant increases were also identified from another two Districts, the Arnisdale (1993 to 1998) and the Kirkaig (1982 to 1994); thus four Districts in the North West Region showed periods of significant increase in sea trout catches.

Five Districts, the Broom, the Carron, the Inver, the Little Loch Broom, and the Sligachan, did not show any statistically significant change in catches across the timeseries. Of the 16 Districts within the North West Region, three Districts, the Kanaird, the Laxford, and the Shiel showed patterns of change identified at the Regional scale (i.e. significant decline in sea trout across the whole timeseries) (Figure 10).

#### Districts in the North Region

The North Region comprises 15 reporting Districts, six of which were included in the cleaned dataset (Figure 11). There were significant differences in total annual sea trout catch between different Districts within the Region (χ^2^_(5)_=372.28, p<0.0001). Statistical difference was identified in 13 pairwise comparisons of Districts (Figure 5f). Within the North Region, the District with the largest mean annual catch was the Hope (806) and the lowest was the Kinloch (14) (Figure 11, Table S2, Table S4).

**Figure 11.**
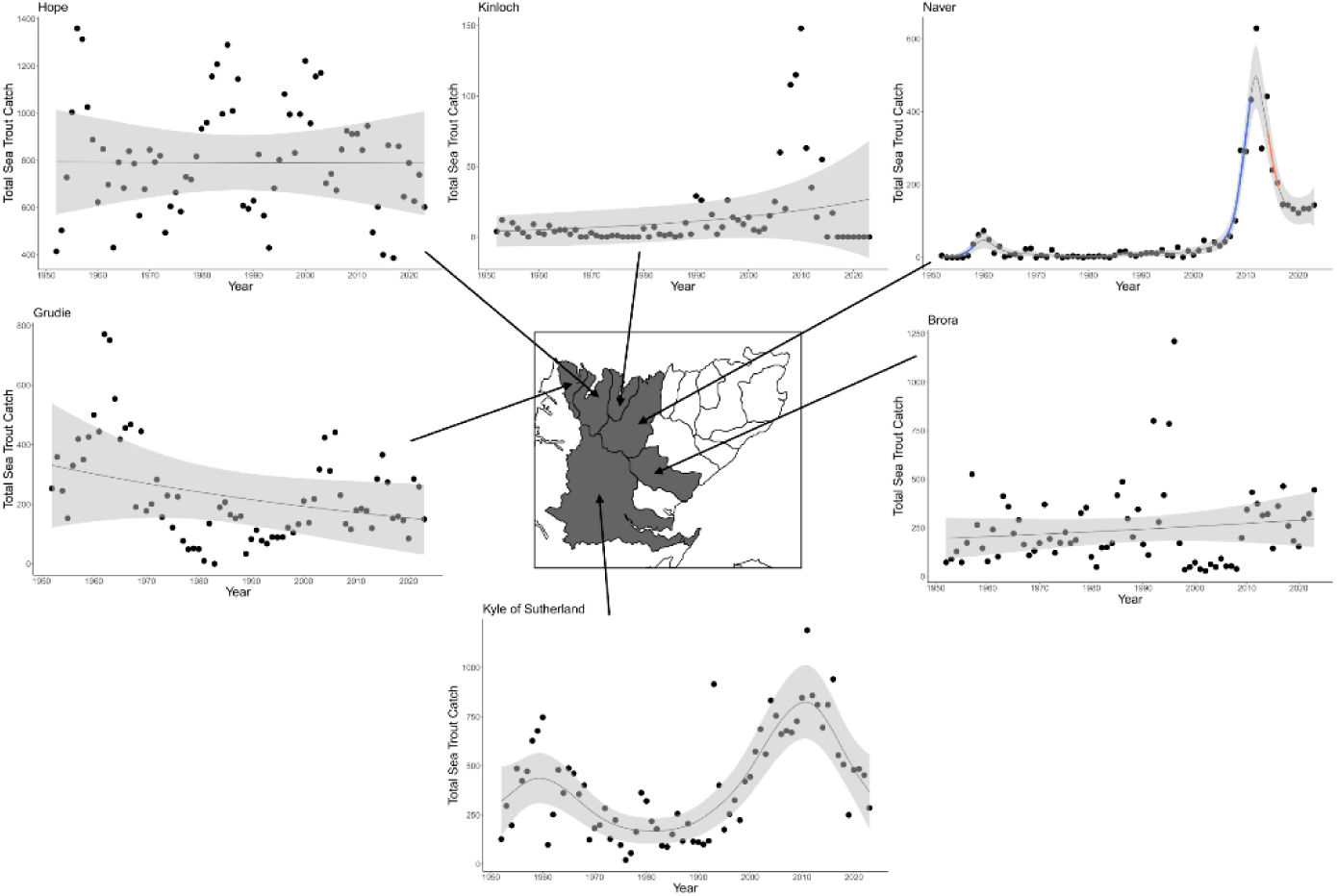
Rod catches of sea trout from each of the six Districts in the North Region (from south to north: the Kyle of Sutherland, Brora, Naver, Hope, Kinloch & Grude) between 1952 and 2023. Solid lines represent the predicted relationship (including the 95% confidence interval as shaded area) from the model (Table S4). Periods of statistically significant increase (blue) and decrease (orange) in rod catches are highlighted. Map redrawn from Scottish Government (2025).

The Naver District was the only one in the North Region to exhibit periods of statistical change (Figure 11, Table 2, Table S2) with a significant increase in sea trout catches (1955 to 1958 and 2005 to 2010) and a period of significant decline (2014 to 2016). The other Districts, the Brora, the Grudie, the Hope, the Kinloch and the Kyle of Sutherland, all showed no statistically significant change in sea trout catches (Figure 11, Table 2). Of the six Districts within the North Region, five Districts (all except the Naver) showed patterns of change identified at the Regional scale (i.e. no significant change across the timeseries).

#### Districts in the Moray Firth Region

The Moray Firth Region comprises nine reporting Districts, seven of which were included in the cleaned dataset (Figure 12). There were significant differences in gross total annual sea trout catch between different Districts within the Region (χ^2^_(6)_ =475.93, p<0.0001). Statistical difference was identified in 19 pairwise comparisons of Districts (Figure 5g). Within the Moray Firth Region, the District with the highest mean annual catch was the Spey (2,904) and the lowest was the Nairn (127) (Table S2, Table S4).

**Figure 12.**
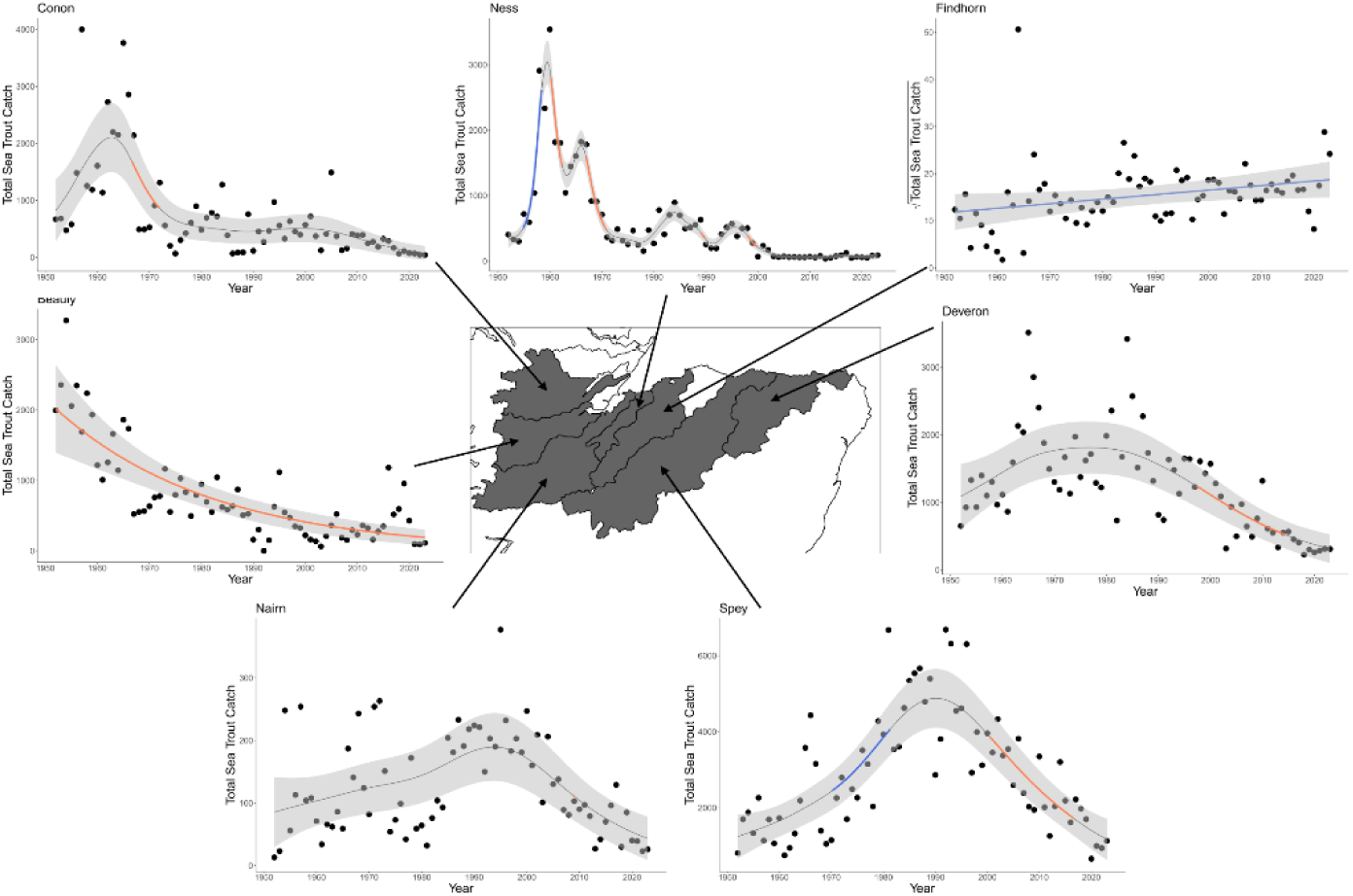
Rod and line catches of sea trout from each of the seven Districts in the Moray Firth region (from west to east: the Nairn, Beauly, Conon, Ness, Findhorn, Spey, Deveron) between 1952 and 2023. Solid lines represent the predicted relationship (including the 95% confidence interval as shaded area) from the model (Table S4). Periods of statistically significant increase (blue) and decrease (orange) in rod catches are highlighted. Map redrawn from Scottish Government (2025).

Eleven statistically significant periods of marked change were identified in seven Districts (Figure 12, Table 2, Table S2, Table S4). Decline in catches were identified in six Districts (Figure 12; Table 2), the Beauly (across the whole timeseries), the Conon (1966 to 1971), the Deveron (1997 to 2014), the Nairn (2008 to 2009), the Ness (1960 to 1962 and 1966 to 1970 and from 1998 to 1999) and the Spey (2001 to 2016). Three Districts had periods of statistically significant increase, the Ness (1954 to 1958) and the Spey (1970 to 1981) and the Findhorn showed a significant (p=0.04) linear increase across the timeseries (Figure 12). Of the seven Districts within the Moray Firth Region, two Districts (the Nairn and the Deveron) showed similar patterns of change (*cf*. Moray Firth significant decline from 1999 to 2013).

#### Districts in the North East Region

The North East Region comprises seven reporting Districts, five of which were included in the cleaned dataset (Figure 13). There were significant differences in total annual sea trout catch between different Districts within the Region (χ^2^_(4)_ =219.32, p<0.0001).

**Figure 13.**
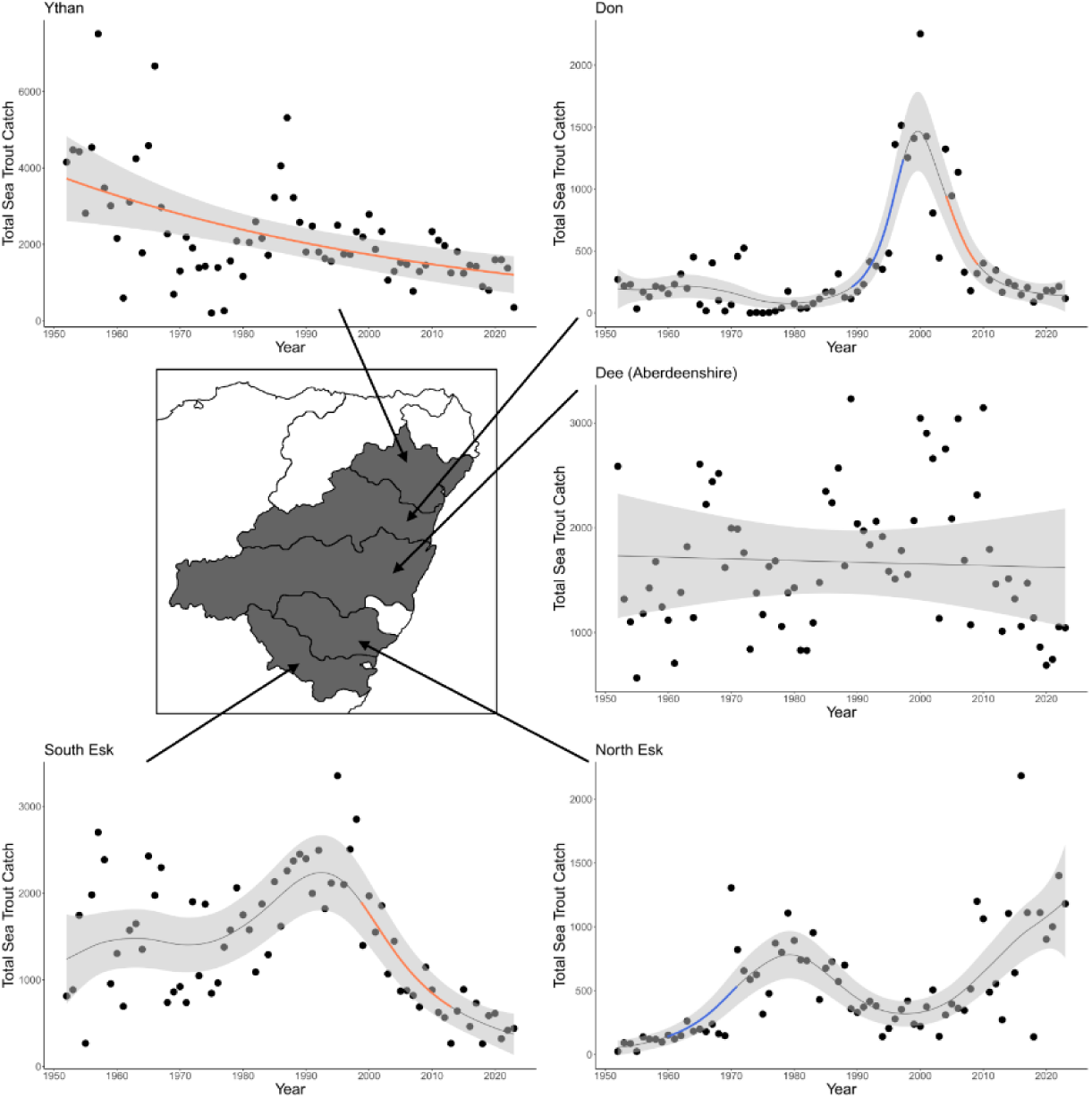
Rod catches of sea trout from each of the five Districts in the North East Region (from north to south: the Ythan, Don, Aberdeenshire Dee, North Esk & South Esk) between 1952 and 2023. Solid lines represent the predicted relationship (including the 95% confidence interval as shaded area) from the model (Table S4). Periods of statistically significant increase (blue) and decrease (orange) in rod catches are highlighted. Map redrawn from Scottish Government (2025).

Statistical difference was identified between eight pairwise comparisons of Districts (Figure 5h). Within the North East Region, the District with the highest mean annual catch was the Ythan (2,239) and the lowest was the Don (356) (Table S2, Table S4).

Five statistically significant periods of marked change were identified in four Districts (Figure 13, Table 2, Table S2, Table S4). Decline in catches were identified in three Districts, the Don (2004 to 2009), the South Esk (1998 to 2013), and the Ythan (across the whole timeseries). There was a period of significant increase in the Don (1989 to 1997) and in the North Esk (1959 to 1970). No significant change in sea trout catch was identified in the Dee (Aberdeenshire). Of the five Districts within the North East Region, one (the Dee (Aberdeenshire)) showed patterns of change identified at the Regional scale (i.e. no significant change across the timeseries).

#### Districts in the East Region

The East Region comprises three reporting Districts, all of which were included in the cleaned dataset (Figure 14). There were significant differences in gross total annual sea trout catch between different Districts within the Region (χ^2^_(2)_ =98.55, p<0.0001). Statistical difference was identified between all pairwise comparisons of Districts (Figure 5i). Within the East Region, the District with the highest mean annual catch was the Tay (1,401) and the lowest was the Forth (493) (Table S2, Table S4).

**Figure 14.**
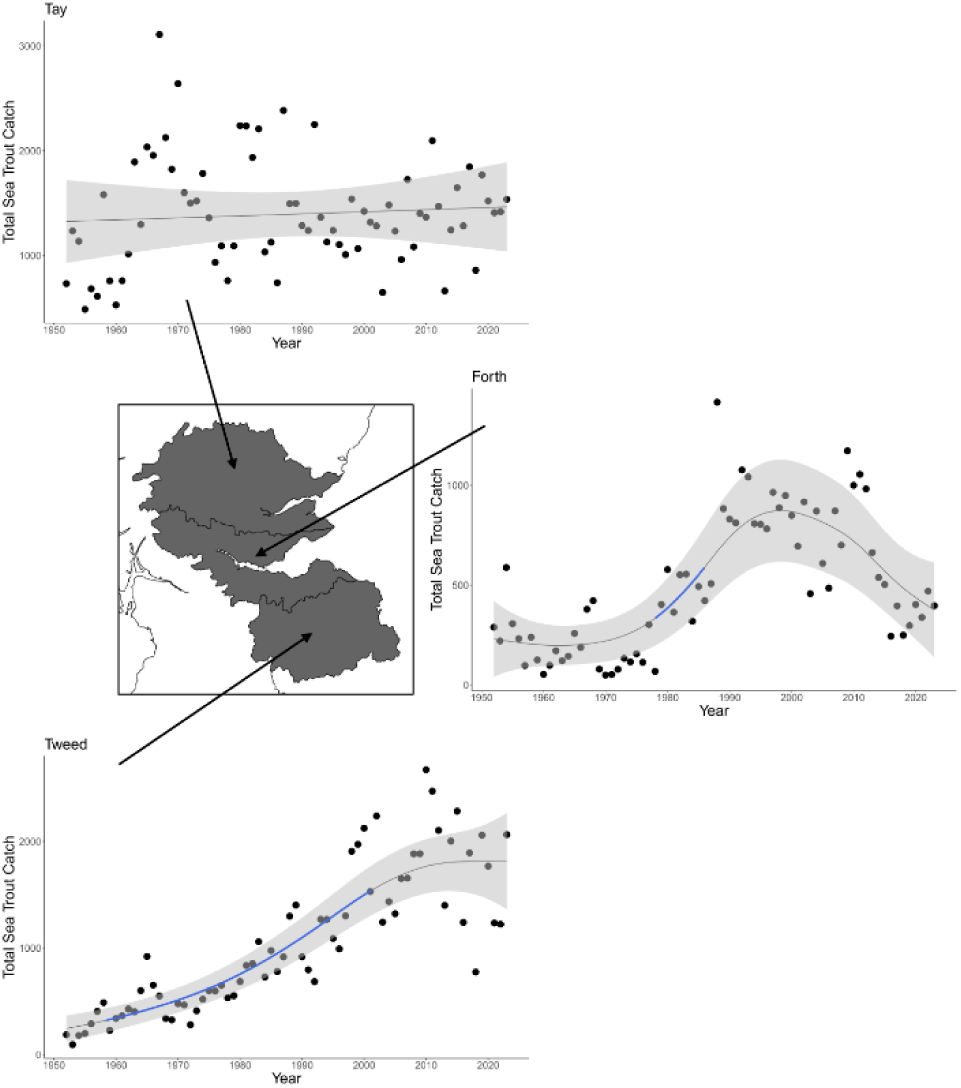
Rod a catches of sea trout from each of the three Districts in the East Region (from north to south: the Tay, Forth & Tweed) between 1952 and 2023. Solid lines represent the predicted relationship (including the 95% confidence interval as shaded area) from the model (Table S4). Periods of statistically significant increase (blue) and decrease (orange) in rod catches are highlighted. Map redrawn from Scottish Government (2025).

Two statistically significant periods of change were identified in two Districts (Figure 14, Table 2, Table S2, Table S4). Increases in catches were identified in two Districts, the Forth (1978 to 1985) and the Tweed (1958 to 2001). No significant change in sea trout catch was identified in the Tay District. Of the three Districts within the East Region, two (the Forth and the Tweed) showed patterns of change identified at the Regional scale (i.e. an increase in catches from 1956 to 1991).

## Discussion

The long-term trends in population size can only be determined through the analysis of information collected over a period of time long enough to subvert the effects of stochastic variation (i.e. process noise) arising from fluctuations in an organism’s biotic and abiotic environment. Here we show that the risk of adequately describing long-term patterns of change may also be affected by the spatial scale over which those trends are examined, with relatively few Districts (n= 13, 20%) and Regions (n=3, 33%) tracking the pattern exhibited at the National scale.

Direct measures of population size for salmonids in general and for sea trout populations in particular in Europe are very limited. Where they do exist they do not cover the broad spatial scale at the spatial density needed for the type of analysis attempted in this study (Shephard *et al*., 2019). To achieve both the high spatial density and scale need for this study we used rod catch returns from fisheries without a measure of fishing effort which have the coverage needed for the questions addressed here. Such data have been shown to be a reasonable proxy for population size in a number of studies. Youngson *et al*. (2002) demonstrated that rod catches provided an accurate depiction of Atlantic salmon abundance trends at an individual catchment level and used the SSSTF dataset (used in the study reported here) to investigate trends of multi-sea winter salmon from 1952 to 1997. Several studies in UK rivers have shown strong relationships between rod catches of Atlantic salmon and numbers from fish counters (Beaumont *et al*., 1991; Crozier & Kennedy, 2001). Adams *et al*. (2022) compared the trends in rod catch data including and excluding a measure of fishing effort and showed that there was no appreciable difference in catch trends between competing models. In addition, comparisons of rod catch data of Atlantic salmon, derived from the SSSTF data, with data from fish counters in 12 Scottish rivers, showed similar trends (Thorley *et al*. 2005). Sea trout rod catch data has been used elsewhere to estimate population size successfully (Millane *et al*., 2017) In the study presented here, rod catches were used not to estimate population size directly but to determine trends in population size over time. Thus we contend that the relative drawbacks resulting from the additional sources of error associated with catch data lacking a measure of effort, is significantly offset by the very considerable temporal and spatial scale of the dataset used here. Therefore, we assume that the temporal and spatial patterns of change that we demonstrate are indicative of underlying changes in sea trout abundance.

At the National scale, the pattern of temporal change is clear and robust; the Scottish sea trout population has declined markedly over the last seven decades, with model estimated population size for the whole of Scotland at the end of the study being 52.8% less than that of the start of the study. Similar patterns of change have been recorded elsewhere (ICES 2013; Harris & Evans, 2017; Fiske et al. 2024). At the Regional level and level of District we also show frequent patterns of decline in the fitted non-linear GAM trend lines, albeit that these patterns are more complex and nuanced. By testing where the GAMs smoother is significantly deviating from a gradient of zero, we were able to identify periods where there was evidence of particularly strong and well defined declines and in some cases increases in catches. By virtue of the fact that the regional and national data comprise the accumulation of data collected at the District level the larger dataset at greater geographic scales makes the detection of trends at broader geographic scales more likely. Thus the periods of change identified at District level should only be regarded as the strongest of such trends.

Where spatial and temporal data for priority species is collected at an appropriate geographical scale, these data can be used to provide population trend information that can be used to inform management and policy decisions at both national and local levels. Funding for national surveys is rarely available however, nor is it often available over the time periods required to develop long-term datasets (Magurran *et al*,. 2010). Direct funding from governments and conservation agency finance can be complex (Cosma *et al*., 2023), often influenced by statutory reporting obligations, as well as social and political factors at both national and local scales. For a number of reasons however, assessment of management needs at a national scale may not fully capture the realistic needs of the species. Here we show that the national pattern of long-term change in the sea trout population is poorly matched at the regional level. The national pattern of population decline was found in only three of nine geographic Regions of Scotland (the three adjacent Regions comprising the Clyde Coast, the West Coast and the North West)). The other Regions exhibited different patterns of population change over time. Therefore it is clear that examination of the long-term patterns of change at national level would result in a different conclusion than that of a similar analysis at regional level. Patterns of sea trout change at the smallest spatial scale (District) also did not closely reflect those observed at the next spatial scale (Region), with only 33% (n=21 of the 64 Districts) of Districts showing a trend similar to that described at the regional level. Comparing patterns of change between the smallest spatial scale with the largest (National) yielded a drop in trend similarity, with 19% (n=12) of Districts exhibiting the national trend of significant decline across the timeseries. Furthermore, and in contrast to this decline, significant increases in sea trout catches were identified in two Districts.

A number of important inferences stem from these results. This study clearly demonstrates that population change in sea trout are primarily governed at the catchment level and less so at regional or national scales. This inference is arguably not at all surprising. The species exhibits very high levels of intraspecific structuring in both expressed phenotype (Ferguson *et al*., 2019b; Klemetsen *et al*., 2003; Koene *et al*., 2020) and genetically (Ferguson, 1989; Verspoor *et al*., 2018) with genetic and phenotypic structuring found between populations over very short geographic distances (Rodger *et al*., 2024, 2021). This is in part, the result of the combination of strong natal site fidelity and of evolutionary responses to subtle environmental differences between juvenile habitats in rivers (King *et al*., 2016; Rodger *et al*., 2024). Despite this, *Salmo trutta* exhibits a continuum of migration strategies, for most populations the rate of straying between spawning sites is low (King *et al*., 2016) but see also (Källo *et al*., 2022 & King et al. 2024) for exceptions. One consequence of the strong structuring of this species is that any environmental change that does occur at a smaller spatial scale is likely to disproportionately affect a smaller number of populations and thus the population dynamics of a proportion of the gene pool (Campbell *et al*., 2017; Koene *et al*., 2024). A second logical inference that the study presented here leads to, is that monitoring population change at the level of a single catchment may not provide an adequate picture of sea trout population change elsewhere. The use of index rivers for monitoring population change in the abundance of Atlantic salmon (ICES 2024; Soulsby *et al*. 2024) and sea trout is widespread (ICES 2013). However, as we show here, it cannot be assumed that the pattern of change detected in these index rivers is reflected in other catchments, regions or nations. Thirdly, the geographic patterns of the changes described at the District level here indicate that even neighbouring catchments can, and do, show very different patterns of population change over time. This points towards a more mosaic pattern of population change, which indicates a relatively low level of confidence in extrapolating patterns from one catchment to even an adjacent catchment. This can have important consequences for identifying the most critical areas for management intervention, thereby ensuring that funding and effort is directed towards those populations that will provide the highest conservation benefit.

Notwithstanding the point about defining units at which management may need to operate to achieve the most effective outcome for sea trout populations with limited resources, it is very clear that when viewed as a national resource, the size of that resource has declined markedly, arguably catastrophically, over the last seven decades with sea trout population size only being around half of the size it was in 1952.

In this study we make no attempt to determine what factors might be driving the changes in sea trout populations describe here. One major potential determinant of the patterns shown here may well be the relative proportion of individuals in any population that adopt an anadromous life history as opposed to remaining to mature in freshwater which has been shown to be influenced by anthropogenic effects (Reed *et al*., 2025). In this study we are not able to disentangle such an effect from any change to the overall population size of both life strategies combined.

## CONTRIBUTIONS

J.A.D. Ideas, data generation, analytical approach, data analysis, manuscript preparation; I.E.M. Ideas, data generation, data analysis, manuscript preparation; A.W.B. Data analysis, manuscript preparation; C.W.B. Ideas, manuscript preparation; JRR, wider context and manuscript preparation; C.E.A. Ideas, data generation, data analysis, manuscript preparation.

## Acknowledgements

This project was part-funded by Grieg Seafood Ltd and the Skye and Lochalsh Rivers Trust. We thank Marine Scotland Science for providing the SSSTF dataset and Marine Scotland Science. Many thanks to G. Ruxton, H. Honkanen, P. Johnson, D. Pascall and M. Meenan for statistical advice and support. We also thank G. Cumming, M. Smith, S. MacIntyre, I. Lindsay and P. Jarosz for project guidance.

## SUPPORTING INFORMATION

## SIGNIFICANCE STATEMENT

In this study we show a complex pattern of both temporal and spatial variation in population size of the anadromous sea trout over a period of seven decades. We demonstrate that national patterns of change are frequently not also seen at smaller spatial scales (regional and district). We show that for this species, assessing the need for conservation management based on national or regional data is likely to miss important trends at smaller spatial scales. We conclude that this may also be true for other species that exhibit high levels of phenotypic and genetic structuring.

## Supplementary material

**Table S1.** The 64 Districts comprising a single catchment or a number of adjacent catchments for where there were consistent data on rod catches of sea trout from Scotland between 1952 and 2023 that were used in this study. Districts are nested into nine broader geographic areas, “Regions” (see Figure 1).

| Region | Catchment | Rivers in the catchment |
| --- | --- | --- |
| Solway | Annan | Annan |
| Solway | Cree | Cree |
| Solway | Luce | Luce |
| Solway | Nith | Nith |
| Solway | Urr | Urr |
| Clyde Coast | Ayr | Ayr |
| Clyde Coast | Carradale | Carradale, Glenlussa |
| Clyde Coast | Clyde | Clyde, Leven, Goil, Endrick |
| Clyde Coast | Doon | Doon |
| Clyde Coast | Eckaig | Eachaig |
| Clyde Coast | Fyne | Kinglas, Cuilarstich, Aray |
| Clyde Coast | Girvan | Girvan |
| Clyde Coast | Irvine | Irvine, Garnock |
| Clyde Coast | Ruel | Ruel |
| Clyde Coast | Stinchar | Stinchar |
| West Coast | Awe | Awe, Etive |
| West Coast | Baa | Ba, Bellart, (Colodoir & Leidle) |
| West Coast | Laggan | (Kintour & Claggain), (Laggan & Sorn) |
| West Coast | Leven | Coe, Leven |
| West Coast | Lochy | Nevis, Lochy |
| West Coast | Nell | Euchar, Nell |
| West Coast | Ormsary | Barr |
| West Coast | Pennygowan | Aros, Forsa |
| North West | Arnisdale | Arnisdale |
| North West | Broom | Broom |
| North West | Carron | Carron |
| North West | Ewe | Ewe |
| North West | Gruinard | Gruinard, Little Gruinard |
| North West | Inver | Inver |
| North West | Kanaird | Kanaird |
| North West | Kirkaig | Kirkaig, (Polly & Osgaig) |
| North West | Laxford | (Laxford & Gleann Dubh), Duartmore |
| North West | Little Loch Broom | Dundonnell |
| North West | Loch Long | (Ling & Elchaig) |
| North West | Moidart | Moidart |
| North West | Morar | Morar |
| North West | Shiel | Shiel, (Achateny & Fascadale) |
| North West | Sligachan | Broadford, Sligachan, Varragill, Lealt, (Brogaig, Stenscholl, Kilmaluag) |
| North West | Snizort | (Camas Fionnairigh (Camasunary) & Scavaig), (Drynoch & Eynort), (Snizort & Ose), (Hinnisdal to Haultin) |
| Outer Hebrides | Creed | Soval estate, Eishken estate, Aline estate, Creed |
| Outer Hebrides | Fincastle | (North Harris SAC), Laxdale, Loch Steisavat |
| Outer Hebrides | Howmore | (Howmaore & Loch Bi), (Kildonan & Bharp) |
| Outer Hebrides | Loch Roag | Langavat SAC, (Caslabhat & Tamanabhaigh), (Mhor a'Ghlinne & Geisiadar), Loch Morsgail sustem, Barvas, Blackwater, Carloway, Forsa |
| North | Brora | Brora, Loth |
| North | Grudie | (KoD - Dionard, Grudie & Daill) |
| North | Hope | Hope, Polla |
| North | Kinloch | Kinloch |
| North | Kyle of Sutherland | Carron, Oykel, Shin, Evelix |
| North | Naver | Naver, Borgie |
| Moray Firth | Beaully | Beaully |
| Moray Firth | Conon | Conon |
| Moray Firth | Deveron | Philorth, Deveron |
| Moray Firth | Findhorn | Findhorn |
| Moray Firth | Nairn | Nairn |
| Moray Firth | Ness | Ness, Moriston |
| Moray Firth | Spey | Spey SAC |
| North East | Dee (Aberdeenshire) | Carron, Cowie, Dee |
| North East | Don | Don |
| North East | North Esk | N Esk |
| North East | South Esk | S Esk |
| North East | Ythan | Ythan |
| East | Forth | Tyne, Almond, Carron, Forth, Teith, Allan, Devon,<br>Leven |
| East | Tay | Eden, Earn, Tay, |
| East | Tweed | River Tweed |

**Table S2.**
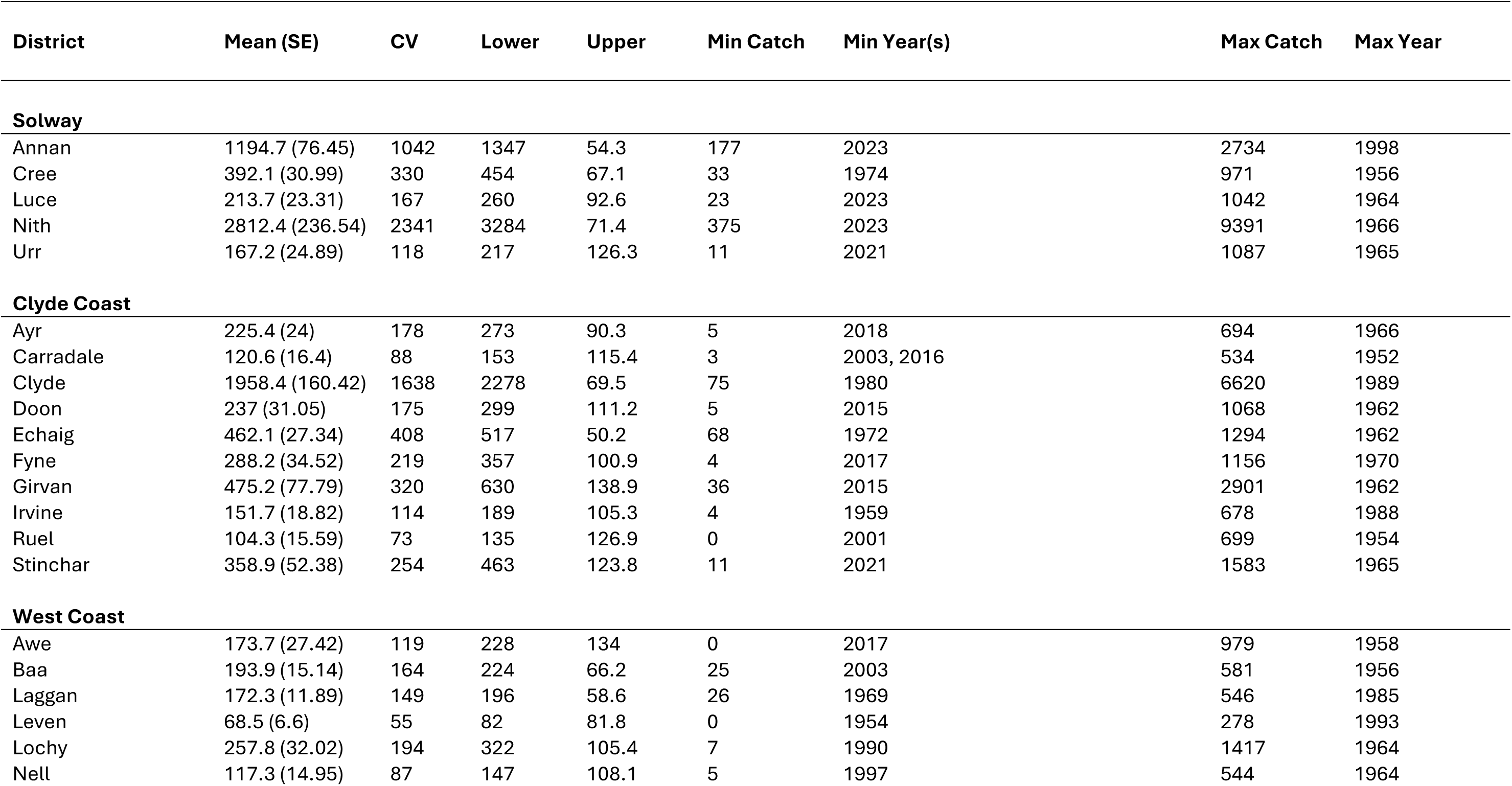
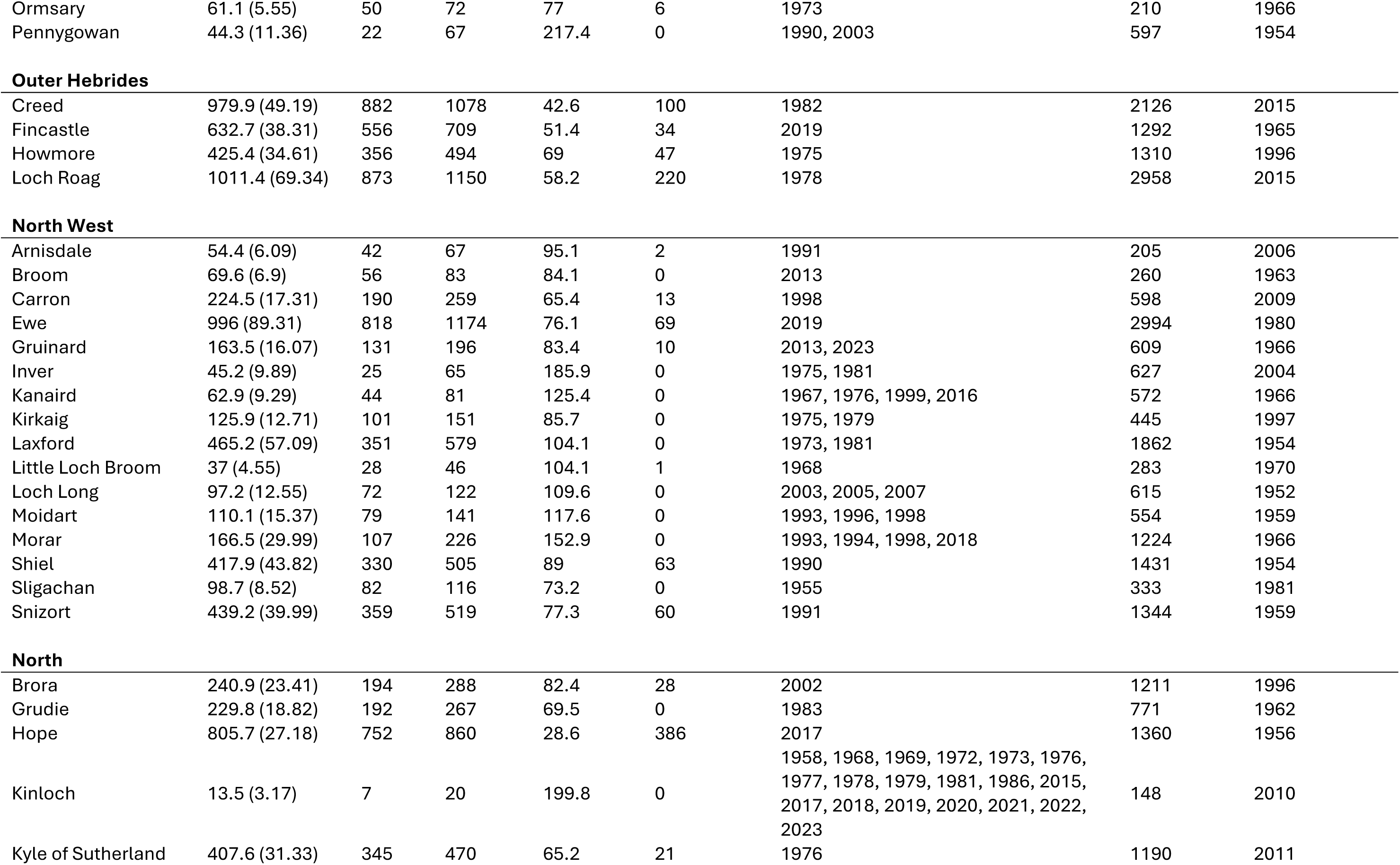
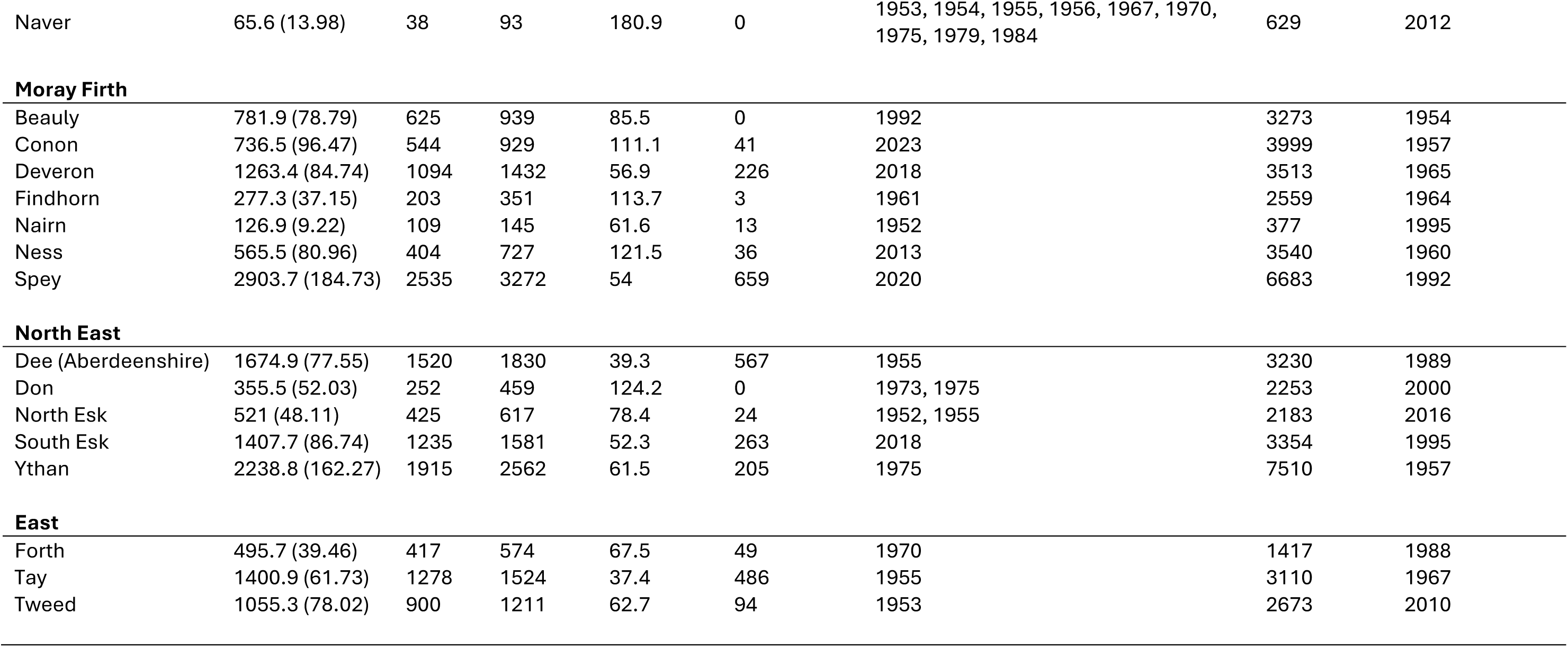
Descriptive data for 64 catchments (Figure 1; Table S1) across 9 Regions (Figure 1) around Scotland over the period of 1952 to 2023. Comprising mean reported sea trout annual rod catch (number, with standard error in parenthesis), coefficient of variation (CV), lower and upper 95% confidence limits, minimum and maximum catch and the year(s) in which the minimum and maximum catches occurred in parenthesis.

| District | Mean (SE) | CV | Lower | Upper | Min Catch | Min Year(s) | Max Catch | Max Year |
| --- | --- | --- | --- | --- | --- | --- | --- | --- |
| **Solway** |  |  |  |  |  |  |  |  |
| Annan | 1194.7 (76.45) | 1042 | 1347 | 54.3 | 177 | 2023 | 2734 | 1998 |
| Cree | 392.1 (30.99) | 330 | 454 | 67.1 | 33 | 1974 | 971 | 1956 |
| Luce | 213.7 (23.31) | 167 | 260 | 92.6 | 23 | 2023 | 1042 | 1964 |
| Nith | 2812.4 (236.54) | 2341 | 3284 | 71.4 | 375 | 2023 | 9391 | 1966 |
| Urr | 167.2 (24.89) | 118 | 217 | 126.3 | 11 | 2021 | 1087 | 1965 |
| **Clyde Coast** |  |  |  |  |  |  |  |  |
| Ayr | 225.4 (24) | 178 | 273 | 90.3 | 5 | 2018 | 694 | 1966 |
| Carradale | 120.6 (16.4) | 88 | 153 | 115.4 | 3 | 2003, 2016 | 534 | 1952 |
| Clyde | 1958.4 (160.42) | 1638 | 2278 | 69.5 | 75 | 1980 | 6620 | 1989 |
| Doon | 237 (31.05) | 175 | 299 | 111.2 | 5 | 2015 | 1068 | 1962 |
| Echaig | 462.1 (27.34) | 408 | 517 | 50.2 | 68 | 1972 | 1294 | 1962 |
| Fyne | 288.2 (34.52) | 219 | 357 | 100.9 | 4 | 2017 | 1156 | 1970 |
| Girvan | 475.2 (77.79) | 320 | 630 | 138.9 | 36 | 2015 | 2901 | 1962 |
| Irvine | 151.7 (18.82) | 114 | 189 | 105.3 | 4 | 1959 | 678 | 1988 |
| Ruel | 104.3 (15.59) | 73 | 135 | 126.9 | 0 | 2001 | 699 | 1954 |
| Stinchar | 358.9 (52.38) | 254 | 463 | 123.8 | 11 | 2021 | 1583 | 1965 |
| **West Coast** |  |  |  |  |  |  |  |  |
| Awe | 173.7 (27.42) | 119 | 228 | 134 | 0 | 2017 | 979 | 1958 |
| Baa | 193.9 (15.14) | 164 | 224 | 66.2 | 25 | 2003 | 581 | 1956 |
| Laggan | 172.3 (11.89) | 149 | 196 | 58.6 | 26 | 1969 | 546 | 1985 |
| Leven | 68.5 (6.6) | 55 | 82 | 81.8 | 0 | 1954 | 278 | 1993 |
| Lochy | 257.8 (32.02) | 194 | 322 | 105.4 | 7 | 1990 | 1417 | 1964 |
| Nell | 117.3 (14.95) | 87 | 147 | 108.1 | 5 | 1997 | 544 | 1964 |
| Ormsary | 61.1 (5.55) | 50 | 72 | 77 | 6 | 1973 | 210 | 1966 |
| Pennygowan | 44.3 (11.36) | 22 | 67 | 217.4 | 0 | 1990, 2003 | 597 | 1954 |
| Outer Hebrides |  |  |  |  |  |  |  |  |
| Creed | 979.9 (49.19) | 882 | 1078 | 42.6 | 100 | 1982 | 2126 | 2015 |
| Fincastle | 632.7 (38.31) | 556 | 709 | 51.4 | 34 | 2019 | 1292 | 1965 |
| Howmore | 425.4 (34.61) | 356 | 494 | 69 | 47 | 1975 | 1310 | 1996 |
| Loch Roag | 1011.4 (69.34) | 873 | 1150 | 58.2 | 220 | 1978 | 2958 | 2015 |
| North West |  |  |  |  |  |  |  |  |
| Arnisdale | 54.4 (6.09) | 42 | 67 | 95.1 | 2 | 1991 | 205 | 2006 |
| Broom | 69.6 (6.9) | 56 | 83 | 84.1 | 0 | 2013 | 260 | 1963 |
| Carron | 224.5 (17.31) | 190 | 259 | 65.4 | 13 | 1998 | 598 | 2009 |
| Ewe | 996 (89.31) | 818 | 1174 | 76.1 | 69 | 2019 | 2994 | 1980 |
| Gruinard | 163.5 (16.07) | 131 | 196 | 83.4 | 10 | 2013, 2023 | 609 | 1966 |
| Inver | 45.2 (9.89) | 25 | 65 | 185.9 | 0 | 1975, 1981 | 627 | 2004 |
| Kanaird | 62.9 (9.29) | 44 | 81 | 125.4 | 0 | 1967, 1976, 1999, 2016 | 572 | 1966 |
| Kirkaig | 125.9 (12.71) | 101 | 151 | 85.7 | 0 | 1975, 1979 | 445 | 1997 |
| Laxford | 465.2 (57.09) | 351 | 579 | 104.1 | 0 | 1973, 1981 | 1862 | 1954 |
| Little Loch Broom | 37 (4.55) | 28 | 46 | 104.1 | 1 | 1968 | 283 | 1970 |
| Loch Long | 97.2 (12.55) | 72 | 122 | 109.6 | 0 | 2003, 2005, 2007 | 615 | 1952 |
| Moidart | 110.1 (15.37) | 79 | 141 | 117.6 | 0 | 1993, 1996, 1998 | 554 | 1959 |
| Morar | 166.5 (29.99) | 107 | 226 | 152.9 | 0 | 1993, 1994, 1998, 2018 | 1224 | 1966 |
| Shiel | 417.9 (43.82) | 330 | 505 | 89 | 63 | 1990 | 1431 | 1954 |
| Sligachan | 98.7 (8.52) | 82 | 116 | 73.2 | 0 | 1955 | 333 | 1981 |
| Snizort | 439.2 (39.99) | 359 | 519 | 77.3 | 60 | 1991 | 1344 | 1959 |
| North |  |  |  |  |  |  |  |  |
| Brora | 240.9 (23.41) | 194 | 288 | 82.4 | 28 | 2002 | 1211 | 1996 |
| Grudie | 229.8 (18.82) | 192 | 267 | 69.5 | 0 | 1983 | 771 | 1962 |
| Hope | 805.7 (27.18) | 752 | 860 | 28.6 | 386 | 2017 | 1360 | 1956 |
| Kinloch | 13.5 (3.17) | 7 | 20 | 199.8 | 0 | 1958, 1968, 1969, 1972, 1973, 1976, 1977, 1978, 1979, 1981, 1986, 2015, 2017, 2018, 2019, 2020, 2021, 2022, 2023 | 148 | 2010 |
| Kyle of Sutherland | 407.6 (31.33) | 345 | 470 | 65.2 | 21 | 1976 | 1190 | 2011 |
| Naver | 65.6 (13.98) | 38 | 93 | 180.9 | 0 | 1953, 1954, 1955, 1956, 1967, 1970,<br>1975, 1979, 1984 | 629 | 2012 |
| **Moray Firth** |  |  |  |  |  |  |  |  |
| Beaully | 781.9 (78.79) | 625 | 939 | 85.5 | 0 | 1992 | 3273 | 1954 |
| Conon | 736.5 (96.47) | 544 | 929 | 111.1 | 41 | 2023 | 3999 | 1957 |
| Deveron | 1263.4 (84.74) | 1094 | 1432 | 56.9 | 226 | 2018 | 3513 | 1965 |
| Findhorn | 277.3 (37.15) | 203 | 351 | 113.7 | 3 | 1961 | 2559 | 1964 |
| Nairn | 126.9 (9.22) | 109 | 145 | 61.6 | 13 | 1952 | 377 | 1995 |
| Ness | 565.5 (80.96) | 404 | 727 | 121.5 | 36 | 2013 | 3540 | 1960 |
| Spey | 2903.7 (184.73) | 2535 | 3272 | 54 | 659 | 2020 | 6683 | 1992 |
| **North East** |  |  |  |  |  |  |  |  |
| Dee (Aberdeenshire) | 1674.9 (77.55) | 1520 | 1830 | 39.3 | 567 | 1955 | 3230 | 1989 |
| Don | 355.5 (52.03) | 252 | 459 | 124.2 | 0 | 1973, 1975 | 2253 | 2000 |
| North Esk | 521 (48.11) | 425 | 617 | 78.4 | 24 | 1952, 1955 | 2183 | 2016 |
| South Esk | 1407.7 (86.74) | 1235 | 1581 | 52.3 | 263 | 2018 | 3354 | 1995 |
| Ythan | 2238.8 (162.27) | 1915 | 2562 | 61.5 | 205 | 1975 | 7510 | 1957 |
| **East** |  |  |  |  |  |  |  |  |
| Forth | 495.7 (39.46) | 417 | 574 | 67.5 | 49 | 1970 | 1417 | 1988 |
| Tay | 1400.9 (61.73) | 1278 | 1524 | 37.4 | 486 | 1955 | 3110 | 1967 |
| Tweed | 1055.3 (78.02) | 900 | 1211 | 62.7 | 94 | 1953 | 2673 | 2010 |

**Table S3.** Periods of statistically significant change in sea trout annual catch in Regions. Numbers reported are model fitted values of sea trout annual catch (modelled catch) including standard error in parenthesis and the lower and upper bounds of the 95% confidence interval for the start year and end year of the period of significant change. For each period of significant change, change in catch is expressed as a percentage of the catch at the start of the period and as a percent change per year.

| Region | Direction | Start |  |  |  | End |  |  |  | % change<br>over the<br>period | %<br>change<br>pa |
| --- | --- | --- | --- | --- | --- | --- | --- | --- | --- | --- | --- |
|  |  | Year | Modelled<br>Catch (SE) | Lower<br>95CI | Upper<br>95CI | Year | Modelled<br>Catch (SE) | Lower<br>95CI | Upper<br>95CI |  |  |
| Solway | Decrease | 1994 | 4229 (660.3) | 5546 | 2911 | 2007 | 2354 (478.7) | 114 | 374 | 55.67 | 4.2 |
| Clyde Coast | Decrease | 1952 | 8405 (1596) | 11588 | 5222 | 2023 | 1832 (534.3) | 26 | 25 | 21.79 | 0.3 |
| West Coast | Decrease | 1952 | 1939 (216.7) | 2371 | 1507 | 2023 | 521 (82.6) | 39 | 206 | 26.9 | 0.3 |
| North West | Decrease | 1952 | 7340 (571.1) | 8479 | 6201 | 2023 | 1368 (164.8) | 69 | 573 | 18.63 | 0.2 |
| Moray Firth | Decrease | 1999 | 6309 (685.3) | 7676 | 4941 | 2012 | 4082 (590.3) | 28 | 120 | 64.7 | 4.9 |
| East | Increase | 1957 | 1803 (218.7) | 2240 | 1367 | 1995 | 3302 (221) | 11 | 22 | 183.09 | 4.8 |

**Table S4.**
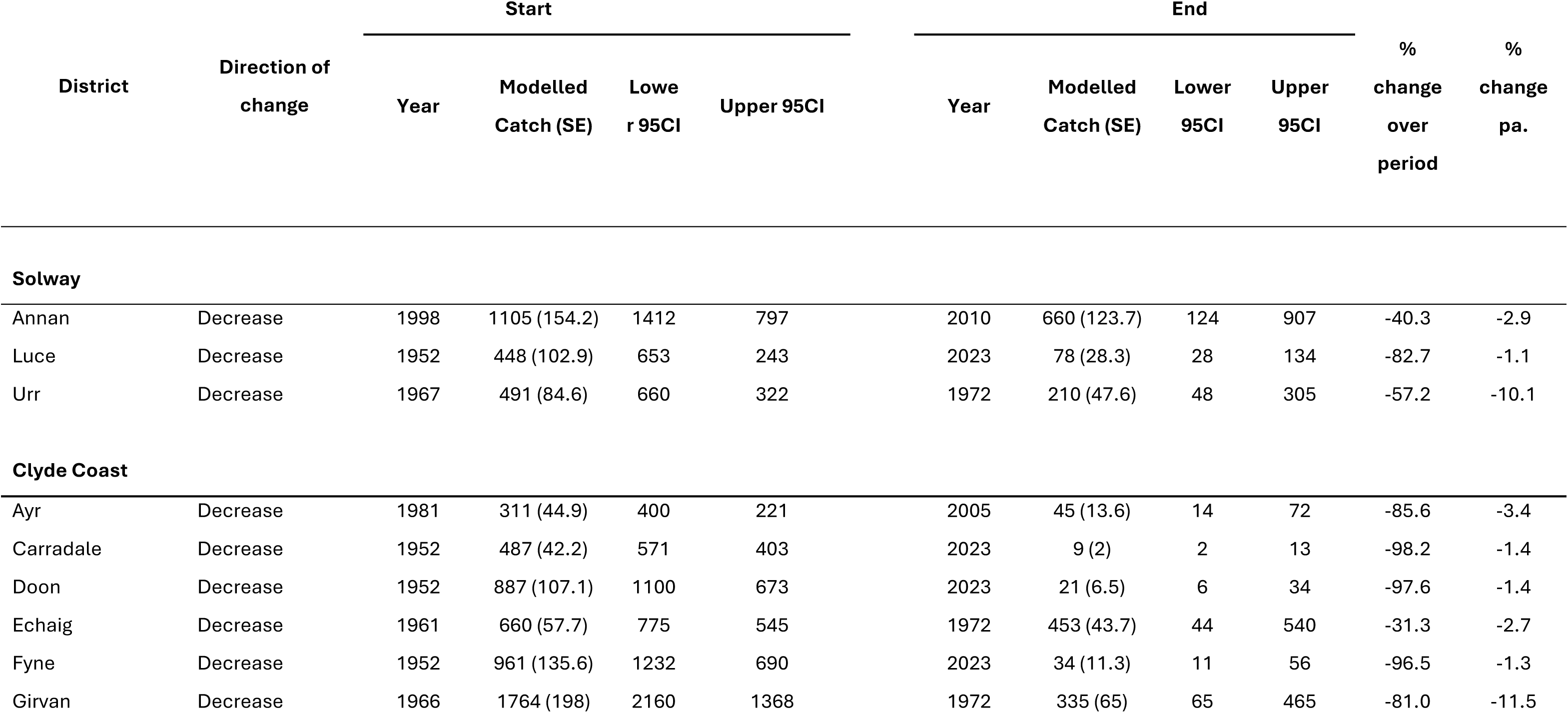
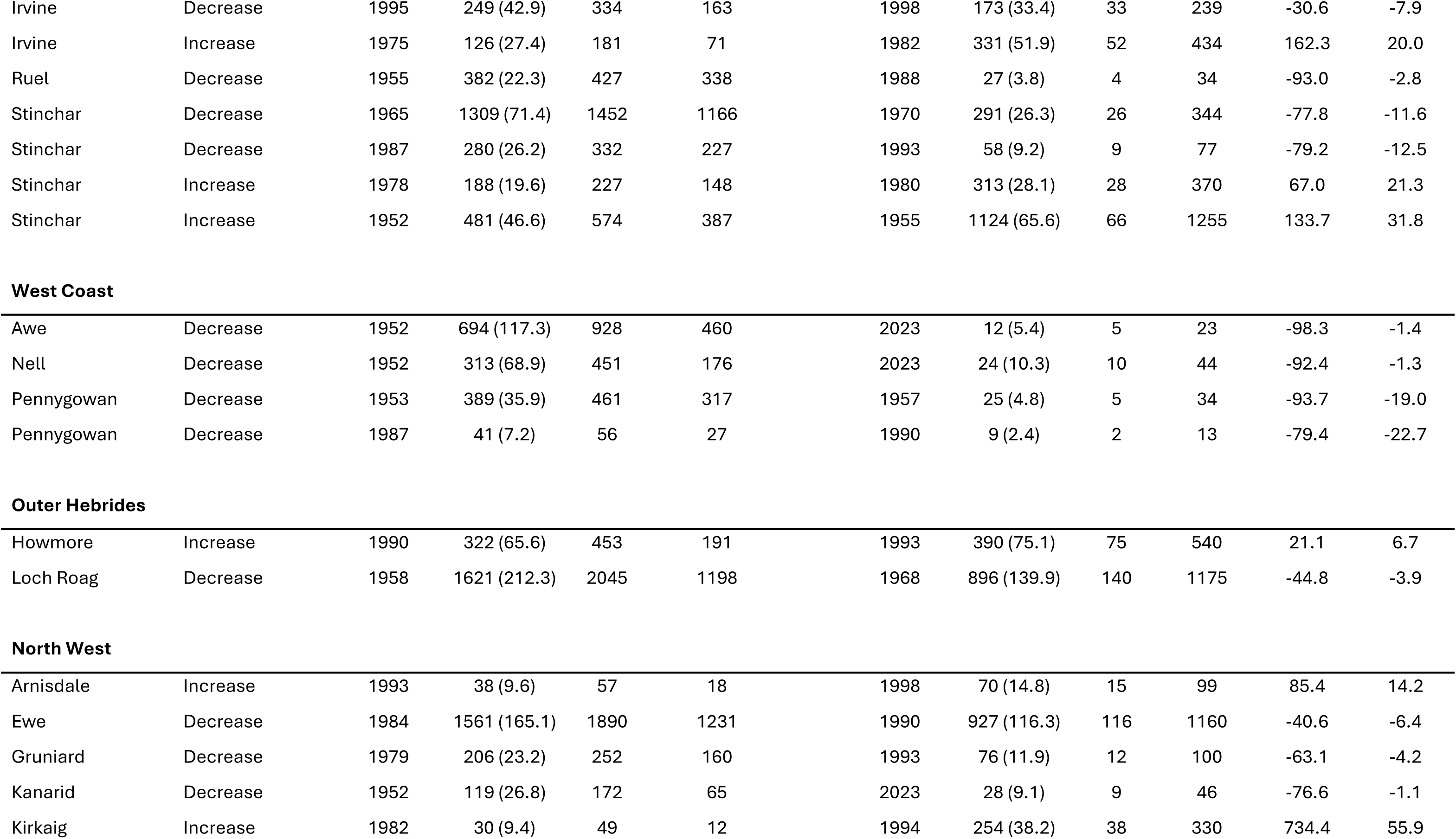
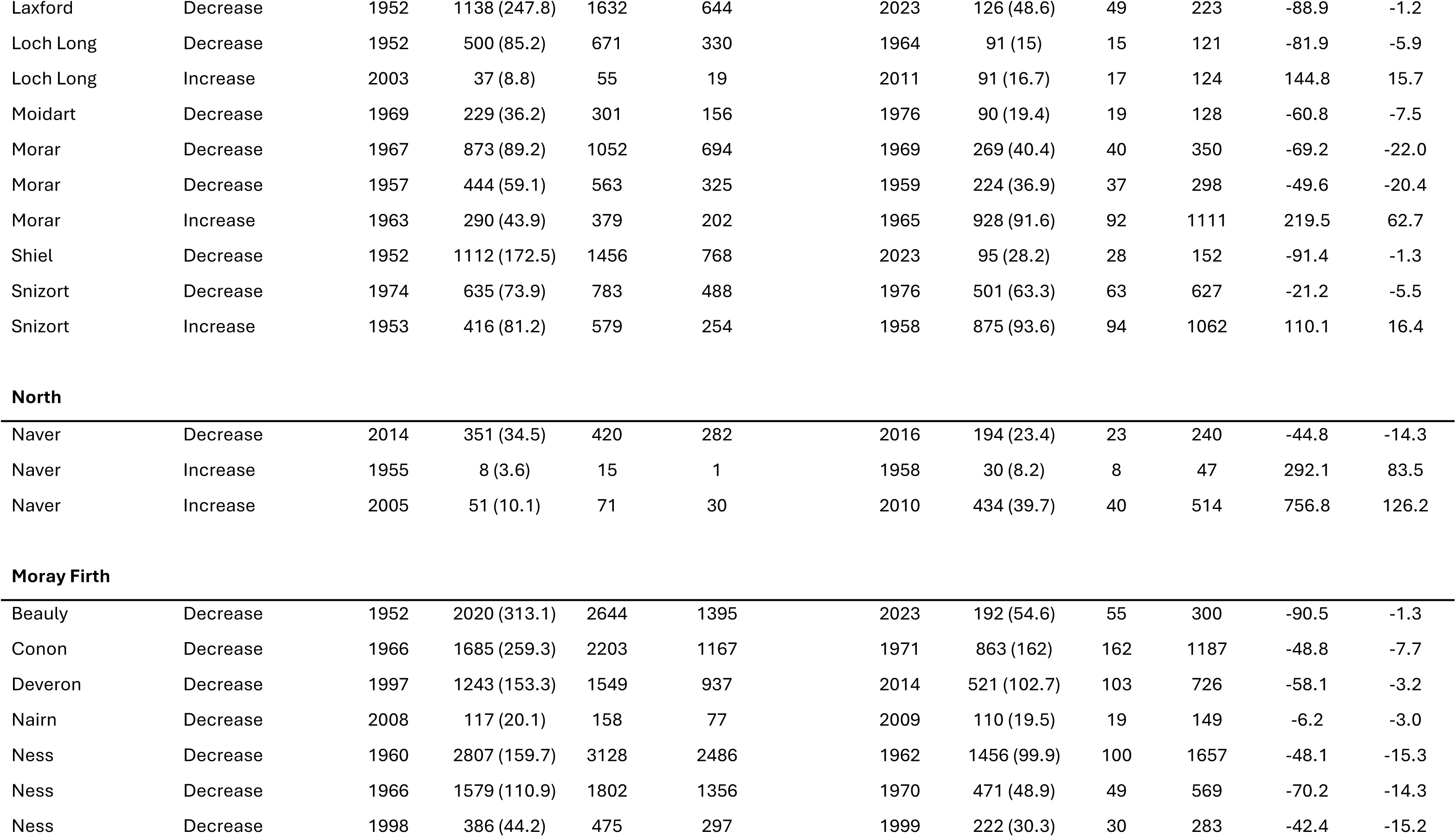
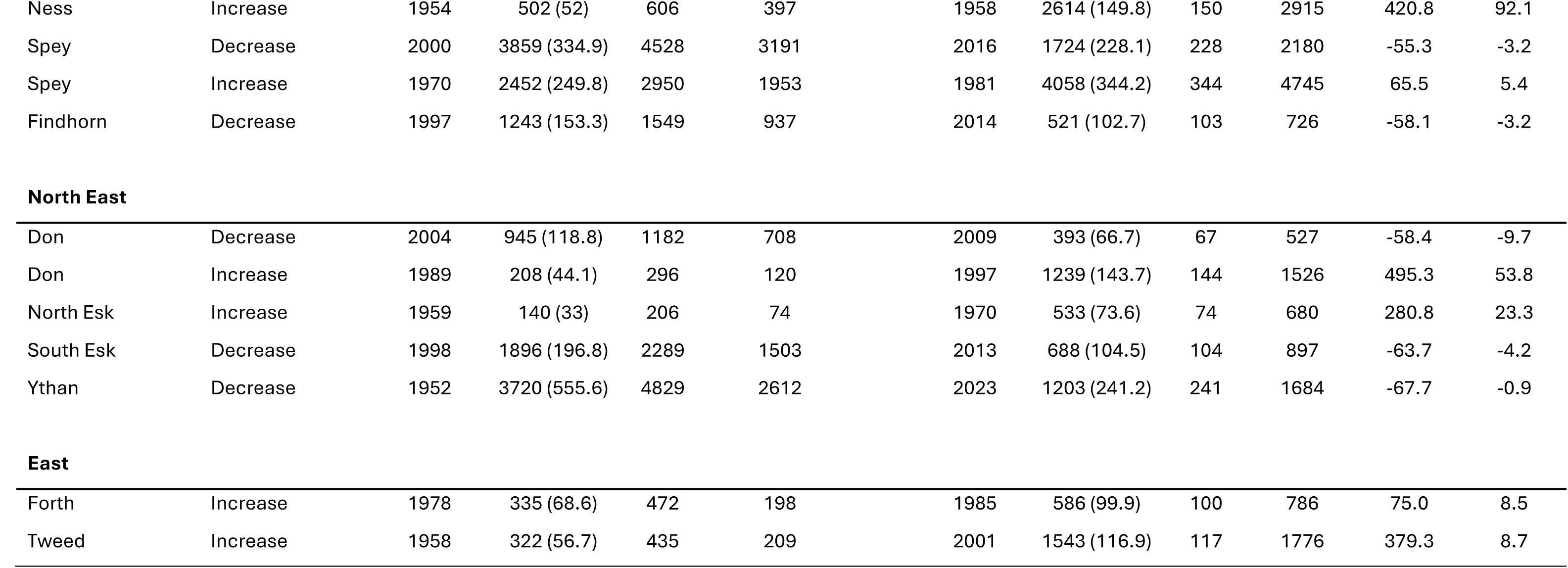
Periods of significant change in sea trout annual catch in Districts. Numbers reported model fitted values of sea trout annual catch (modelled catch) including standard error in parenthesis and the lower and upper bounds of the 95% confidence interval for the start year and end year of the period of significant change. For each period of significant change, change in catch is expressed as a percentage of the catch at the start of the period and as a percent change per year. Districts are grouped by Region then listed alphabetically.

| District | Direction of change | Start |  |  |  | End |  |  |  | % change over period | % change pa. |
| --- | --- | --- | --- | --- | --- | --- | --- | --- | --- | --- | --- |
|  |  | Year | Modelled Catch (SE) | Lower 95CI | Upper 95CI | Year | Modelled Catch (SE) | Lower 95CI | Upper 95CI |  |  |
| Solway |  |  |  |  |  |  |  |  |  |  |  |
| Annan | Decrease | 1998 | 1105 (154.2) | 1412 | 797 | 2010 | 660 (123.7) | 124 | 907 | -40.3 | -2.9 |
| Luce | Decrease | 1952 | 448 (102.9) | 653 | 243 | 2023 | 78 (28.3) | 28 | 134 | -82.7 | -1.1 |
| Urr | Decrease | 1967 | 491 (84.6) | 660 | 322 | 1972 | 210 (47.6) | 48 | 305 | -57.2 | -10.1 |
| Clyde Coast |  |  |  |  |  |  |  |  |  |  |  |
| Ayr | Decrease | 1981 | 311 (44.9) | 400 | 221 | 2005 | 45 (13.6) | 14 | 72 | -85.6 | -3.4 |
| Carradale | Decrease | 1952 | 487 (42.2) | 571 | 403 | 2023 | 9 (2) | 2 | 13 | -98.2 | -1.4 |
| Doon | Decrease | 1952 | 887 (107.1) | 1100 | 673 | 2023 | 21 (6.5) | 6 | 34 | -97.6 | -1.4 |
| Echaig | Decrease | 1961 | 660 (57.7) | 775 | 545 | 1972 | 453 (43.7) | 44 | 540 | -31.3 | -2.7 |
| Fyne | Decrease | 1952 | 961 (135.6) | 1232 | 690 | 2023 | 34 (11.3) | 11 | 56 | -96.5 | -1.3 |
| Girvan | Decrease | 1966 | 1764 (198) | 2160 | 1368 | 1972 | 335 (65) | 65 | 465 | -81.0 | -11.5 |
| Irvine | Decrease | 1995 | 249 (42.9) | 334 | 163 | 1998 | 173 (33.4) | 33 | 239 | -30.6 | -7.9 |
| Irvine | Increase | 1975 | 126 (27.4) | 181 | 71 | 1982 | 331 (51.9) | 52 | 434 | 162.3 | 20.0 |
| Ruel | Decrease | 1955 | 382 (22.3) | 427 | 338 | 1988 | 27 (3.8) | 4 | 34 | -93.0 | -2.8 |
| Stinchar | Decrease | 1965 | 1309 (71.4) | 1452 | 1166 | 1970 | 291 (26.3) | 26 | 344 | -77.8 | -11.6 |
| Stinchar | Decrease | 1987 | 280 (26.2) | 332 | 227 | 1993 | 58 (9.2) | 9 | 77 | -79.2 | -12.5 |
| Stinchar | Increase | 1978 | 188 (19.6) | 227 | 148 | 1980 | 313 (28.1) | 28 | 370 | 67.0 | 21.3 |
| Stinchar | Increase | 1952 | 481 (46.6) | 574 | 387 | 1955 | 1124 (65.6) | 66 | 1255 | 133.7 | 31.8 |
| West Coast |  |  |  |  |  |  |  |  |  |  |  |
| Awe | Decrease | 1952 | 694 (117.3) | 928 | 460 | 2023 | 12 (5.4) | 5 | 23 | -98.3 | -1.4 |
| Nell | Decrease | 1952 | 313 (68.9) | 451 | 176 | 2023 | 24 (10.3) | 10 | 44 | -92.4 | -1.3 |
| Pennygowan | Decrease | 1953 | 389 (35.9) | 461 | 317 | 1957 | 25 (4.8) | 5 | 34 | -93.7 | -19.0 |
| Pennygowan | Decrease | 1987 | 41 (7.2) | 56 | 27 | 1990 | 9 (2.4) | 2 | 13 | -79.4 | -22.7 |
| Outer Hebrides |  |  |  |  |  |  |  |  |  |  |  |
| Howmore | Increase | 1990 | 322 (65.6) | 453 | 191 | 1993 | 390 (75.1) | 75 | 540 | 21.1 | 6.7 |
| Loch Roag | Decrease | 1958 | 1621 (212.3) | 2045 | 1198 | 1968 | 896 (139.9) | 140 | 1175 | -44.8 | -3.9 |
| North West |  |  |  |  |  |  |  |  |  |  |  |
| Arnisdale | Increase | 1993 | 38 (9.6) | 57 | 18 | 1998 | 70 (14.8) | 15 | 99 | 85.4 | 14.2 |
| Ewe | Decrease | 1984 | 1561 (165.1) | 1890 | 1231 | 1990 | 927 (116.3) | 116 | 1160 | -40.6 | -6.4 |
| Gruniard | Decrease | 1979 | 206 (23.2) | 252 | 160 | 1993 | 76 (11.9) | 12 | 100 | -63.1 | -4.2 |
| Kanarid | Decrease | 1952 | 119 (26.8) | 172 | 65 | 2023 | 28 (9.1) | 9 | 46 | -76.6 | -1.1 |
| Kirkaig | Increase | 1982 | 30 (9.4) | 49 | 12 | 1994 | 254 (38.2) | 38 | 330 | 734.4 | 55.9 |
| Laxford | Decrease | 1952 | 1138 (247.8) | 1632 | 644 | 2023 | 126 (48.6) | 49 | 223 | -88.9 | -1.2 |
| Loch Long | Decrease | 1952 | 500 (85.2) | 671 | 330 | 1964 | 91 (15) | 15 | 121 | -81.9 | -5.9 |
| Loch Long | Increase | 2003 | 37 (8.8) | 55 | 19 | 2011 | 91 (16.7) | 17 | 124 | 144.8 | 15.7 |
| Moidart | Decrease | 1969 | 229 (36.2) | 301 | 156 | 1976 | 90 (19.4) | 19 | 128 | -60.8 | -7.5 |
| Morar | Decrease | 1967 | 873 (89.2) | 1052 | 694 | 1969 | 269 (40.4) | 40 | 350 | -69.2 | -22.0 |
| Morar | Decrease | 1957 | 444 (59.1) | 563 | 325 | 1959 | 224 (36.9) | 37 | 298 | -49.6 | -20.4 |
| Morar | Increase | 1963 | 290 (43.9) | 379 | 202 | 1965 | 928 (91.6) | 92 | 1111 | 219.5 | 62.7 |
| Shiel | Decrease | 1952 | 1112 (172.5) | 1456 | 768 | 2023 | 95 (28.2) | 28 | 152 | -91.4 | -1.3 |
| Snizort | Decrease | 1974 | 635 (73.9) | 783 | 488 | 1976 | 501 (63.3) | 63 | 627 | -21.2 | -5.5 |
| Snizort | Increase | 1953 | 416 (81.2) | 579 | 254 | 1958 | 875 (93.6) | 94 | 1062 | 110.1 | 16.4 |
| North |  |  |  |  |  |  |  |  |  |  |  |
| Naver | Decrease | 2014 | 351 (34.5) | 420 | 282 | 2016 | 194 (23.4) | 23 | 240 | -44.8 | -14.3 |
| Naver | Increase | 1955 | 8 (3.6) | 15 | 1 | 1958 | 30 (8.2) | 8 | 47 | 292.1 | 83.5 |
| Naver | Increase | 2005 | 51 (10.1) | 71 | 30 | 2010 | 434 (39.7) | 40 | 514 | 756.8 | 126.2 |
| Moray Firth |  |  |  |  |  |  |  |  |  |  |  |
| Beauly | Decrease | 1952 | 2020 (313.1) | 2644 | 1395 | 2023 | 192 (54.6) | 55 | 300 | -90.5 | -1.3 |
| Conon | Decrease | 1966 | 1685 (259.3) | 2203 | 1167 | 1971 | 863 (162) | 162 | 1187 | -48.8 | -7.7 |
| Deveron | Decrease | 1997 | 1243 (153.3) | 1549 | 937 | 2014 | 521 (102.7) | 103 | 726 | -58.1 | -3.2 |
| Nairn | Decrease | 2008 | 117 (20.1) | 158 | 77 | 2009 | 110 (19.5) | 19 | 149 | -6.2 | -3.0 |
| Ness | Decrease | 1960 | 2807 (159.7) | 3128 | 2486 | 1962 | 1456 (99.9) | 100 | 1657 | -48.1 | -15.3 |
| Ness | Decrease | 1966 | 1579 (110.9) | 1802 | 1356 | 1970 | 471 (48.9) | 49 | 569 | -70.2 | -14.3 |
| Ness | Decrease | 1998 | 386 (44.2) | 475 | 297 | 1999 | 222 (30.3) | 30 | 283 | -42.4 | -15.2 |
| Ness | Increase | 1954 | 502 (52) | 606 | 397 | 1958 | 2614 (149.8) | 150 | 2915 | 420.8 | 92.1 |
| Spey | Decrease | 2000 | 3859 (334.9) | 4528 | 3191 | 2016 | 1724 (228.1) | 228 | 2180 | -55.3 | -3.2 |
| Spey | Increase | 1970 | 2452 (249.8) | 2950 | 1953 | 1981 | 4058 (344.2) | 344 | 4745 | 65.5 | 5.4 |
| Findhorn | Decrease | 1997 | 1243 (153.3) | 1549 | 937 | 2014 | 521 (102.7) | 103 | 726 | -58.1 | -3.2 |
| **North East** |  |  |  |  |  |  |  |  |  |  |  |
| Don | Decrease | 2004 | 945 (118.8) | 1182 | 708 | 2009 | 393 (66.7) | 67 | 527 | -58.4 | -9.7 |
| Don | Increase | 1989 | 208 (44.1) | 296 | 120 | 1997 | 1239 (143.7) | 144 | 1526 | 495.3 | 53.8 |
| North Esk | Increase | 1959 | 140 (33) | 206 | 74 | 1970 | 533 (73.6) | 74 | 680 | 280.8 | 23.3 |
| South Esk | Decrease | 1998 | 1896 (196.8) | 2289 | 1503 | 2013 | 688 (104.5) | 104 | 897 | -63.7 | -4.2 |
| Ythan | Decrease | 1952 | 3720 (555.6) | 4829 | 2612 | 2023 | 1203 (241.2) | 241 | 1684 | -67.7 | -0.9 |
| **East** |  |  |  |  |  |  |  |  |  |  |  |
| Forth | Increase | 1978 | 335 (68.6) | 472 | 198 | 1985 | 586 (99.9) | 100 | 786 | 75.0 | 8.5 |
| Tweed | Increase | 1958 | 322 (56.7) | 435 | 209 | 2001 | 1543 (116.9) | 117 | 1776 | 379.3 | 8.7 |

**Figure S1.**
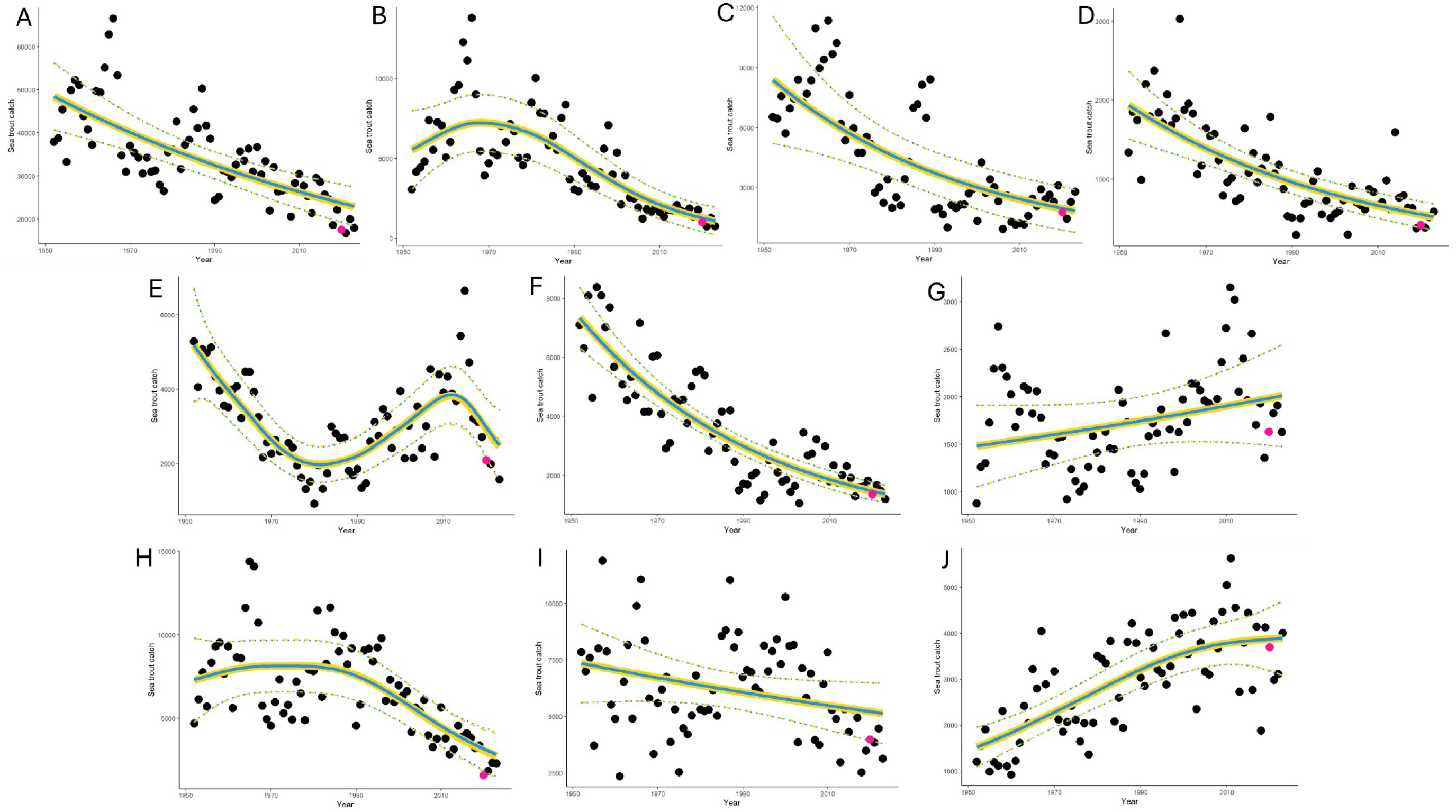
Sensitivity analysis assessing the inclusion of data from 2020 (the year when access to angling may have been affected by restrictions resulting from the global COVID virus pandemic). For national, regional and district catch data. A = national, B = Solway, C = Clyde Coast, D = West Coast, E = Outer Hebrides, F = North West, G = North, H = Moray Firth, I = North East, J = East.

## Notes

### Competing Interest Statement

The authors have declared no competing interest.

